# Patient-Derived Inner Ear Organoids as a Disease Modeling and Therapy Validation Platform For Hereditary Inner Ear Disorders

**DOI:** 10.64898/2026.07.14.738390

**Authors:** Esther Fousert, Winnie M.C. van den Boogaard, Amy W.A. Lucassen, Shantha D. Udayappan, Jaap Oostrik, Peter Paul G. van Benthem, Hannie Kremer, Erwin van Wijk, Wouter H. van der Valk, Erik de Vrieze, Heiko Locher

## Abstract

**Background:** Hereditary inner ear disorders comprise a highly heterogenous group of disorders and are a major cause of hearing and vestibular dysfunction. Despite advances in genetic diagnosis, the development of precision therapies has been limited by the lack of relevant and scalable human model systems that can accommodate the wide spectrum of disease-causing variants and support the evaluation of therapeutic interventions. We established patient-derived inner ear organoids (IEOs) as a platform to assess antisense oligonucleotide (ASO)-based therapeutic strategies for hereditary hearing loss.

**Methods:** Two representative genetic models were selected: recessive syndromic Usher syndrome type IIa (*USH2A*) and dominant non-syndromic DFNA9 (*COCH*). Human induced pluripotent stem cells (iPSCs) were generated from a patient carrying a homozygous pathogenic *USH2A* variant and a patient carrying a frequently occurring pathogenic *COCH* variant. In parallel, isogenic iPSC lines were created by introducing the same disease-causing variants into a healthy donor background. Following differentiation into IEOs, disease-associated transcript expression was evaluated. Splice-switching and RNase H1-mediated gapmer ASOs were assessed for target engagement. ASO biodistribution and cellular uptake was also examined in both IEOs and adult human vestibular tissue.

**Results:** Patient-derived and isogenic iPSCs were successfully differentiated into IEOs that recapitulated disease-associated transcript expression. ASOs showed efficient uptake into disease-relevant cell populations in both IEOs and adult human vestibular tissue. In *USH2A*-variant IEOs, splice-switching ASO treatment corrected aberrant splicing. In *COCH*-variant IEOs, gapmer ASO treatment reduced total *COCH* transcript levels, achieving up to 75% knockdown in patient-derived IEOs.

**Conclusions:** Patient-derived and isogenic variant IEOs provide a versatile and scalable human platform for evaluating ASO therapies for hereditary hearing loss. Their adaptability to diverse genetic variants, inheritance patterns, and ASO modalities makes them well suited to address the genetic heterogeneity of hereditary inner ear diseases and establishes IEOs as a broadly applicable preclinical model for rare hereditary inner ear diseases.

## Introduction

Hereditary hearing loss is the most prevalent and genetically heterogenous sensory disorder worldwide (1,2). Pathogenic variants in numerous genes give rise to a wide spectrum of audiological and vestibular phenotypes, ranging from congenital anatomical malformations to dysfunction of specific proteins and respective functional loss (3–5). Despite advancements in genetic diagnosis (6), current treatment strategies remain focused on the use of hearing aids and ultimately cochlear implants. Although these devices improve auditory perception, this is not achieved over all frequences, nor do they address vestibular dysfunctions or halt degeneration in progressive inner ear disorders. Recent progress in molecular therapies, shown by adeno-associated virus (AAV)-mediated gene therapy for *OTOF*-related deafness (7–10), shows that treating the underlying cause of hereditary hearing loss with genetic strategies is possible (11). However, such approaches remain applicable to small patient cohorts and do not yet address the high diversity of genetic variants associated with hereditary inner ear disease (12,13).

A major barrier to the development of disease-modifying strategies is the limited availability of representative preclinical models of human hereditary inner ear diseases (14). Animal models have provided fundamental insights into how the inner ear develops and functions. However, translation to humans remains challenging due to interspecies differences in gene sequence, expression and regulation as well as inner ear anatomy (15,16). In addition, the large number of unique and often (ultra)rare pathogenic variants underlying hereditary hearing and vestibular disorders emphasize the need for a preclinical model that is flexible and scalable to unravel pathogenic mechanisms of disease and evaluate personalized therapeutic interventions (17).

Human inner ear organoids (IEOs) derived from induced pluripotent stem cells (hiPSCs) are a promising *in vitro* model of the human inner ear (18–20). These three-dimensional cultures recapitulate key aspects of early inner ear development and contain various cell types, including sensory epithelia with hair cells and supporting cells, as well as neuronal innervation and otic mesenchymal cells (19). To date, most studies have focused on IEOs derived from healthy donor hiPSCs (21–23), or from genetically engineered hiPSC lines in which known pathogenic variants were introduced (24,25) to study development or pathology-related processes. Patient-derived IEOs provide a model with the genetic context of a patient and circumvent the inaccessibility of inner ear tissue, making it a suitable platform for disease modelling and therapeutic testing that can be easily adapted depending on the disease context. Recently, IEOs have indeed been proposed as a platform for testing of disease-modifying interventions (26). However, studies showing the successful use of patient-derived IEOs to investigate the efficacy of such treatments have not been published before.

In parallel with the advances in organoid technology, antisense oligonucleotides (ASOs) have emerged as a versatile and clinically validated therapeutic strategy for hereditary diseases (27–30). ASOs are short synthetic nucleic acids that bind target (pre)mRNA sequences and modulate the expression of transcripts by interfering with splicing or expression (31). Through these mechanisms, ASOs can selectively suppress mutant alleles or restore normal gene expression by correcting pre-mRNA splicing. ASOs can be rapidly adapted to target diverse genes and variants, without the need for viral vectors for delivery. This flexibility makes ASOs suited for distinct pathogenic variants related to hereditary hearing loss (29,32–36). Several ASO-based therapies have already progressed from preclinical development to clinical use, underscoring their translational potential (30).

In this study, we generated patient-derived IEOs to investigate their suitability as a preclinical model for ASO testing for hereditary hearing loss, with confirmation of target engagement in primary human inner ear tissue. To assess the generalizability of our approach across different forms of hereditary hearing loss, we selected two genes representing distinct cell populations, disease mechanisms and clinical phenotypes: *USH2A* and *COCH*. Mutations in *USH2A* are a major cause of Usher Syndrome Type II in which congenital hearing loss is; accompanied with retinitis pigmentosa in childhood or adolescence (37,38). Mutations in *COCH* are associated with non-syndromic hearing loss type DFNA9 characterized by progressive, late-onset vestibular and auditory dysfunction (39,40). With this selection we cover recessive and dominant inheritance, syndromic and non-syndromic disease, early- and late-onset hearing loss, and two complementary ASO modalities: splice-switching and RNase H1-mediated gapmer ASOs. Using these complementary disease models, we assessed the utility of patient-derived IEOs as a platform for evaluating gene-specific ASO therapies. Our findings position patient-derived IEOs as a tool for bridging the gap between genetic diagnosis and the development of molecular treatments for hereditary inner ear diseases.

## Results

### Patient-derived inner ear organoids establish a platform for modeling hereditary hearing loss

From the set of known hearing loss-associated genes (Figure 1a), we selected *USH2A* and *COCH* to model two hereditary inner ear diseases with a distinct pathophysiology (Figure 1b). Using a recent single-nucleus transcriptomic atlas of the developing human fetal inner ear (41), we confirmed the expression of *USH2A* transcripts in hair cells and *COCH* expression in the embryonic precursor cells of the adult inner ear fibrocytes, the otic mesenchymal cells (OMCs; Figure S1), consistent with previous studies (42,43). We next evaluated *USH2A* and *COCH* expression in IEOs between days 75-110 of differentiation (D75-110; (19)), a stage at which IEOs contain sensory epithelia with hair cells and OMCs. Similar to the expression patterns observed in the fetal human inner ear, *USH2A* expression was restricted to hair cell populations in the IEOs, whereas *COCH* showed moderate expression in the OMC population (Figure 1c). Together, these findings encourage further investigation into IEOs as human models for studying *USH2A* and *COCH* variants and evaluating therapeutic strategies in a disease-relevant context.

**Figure 1.**
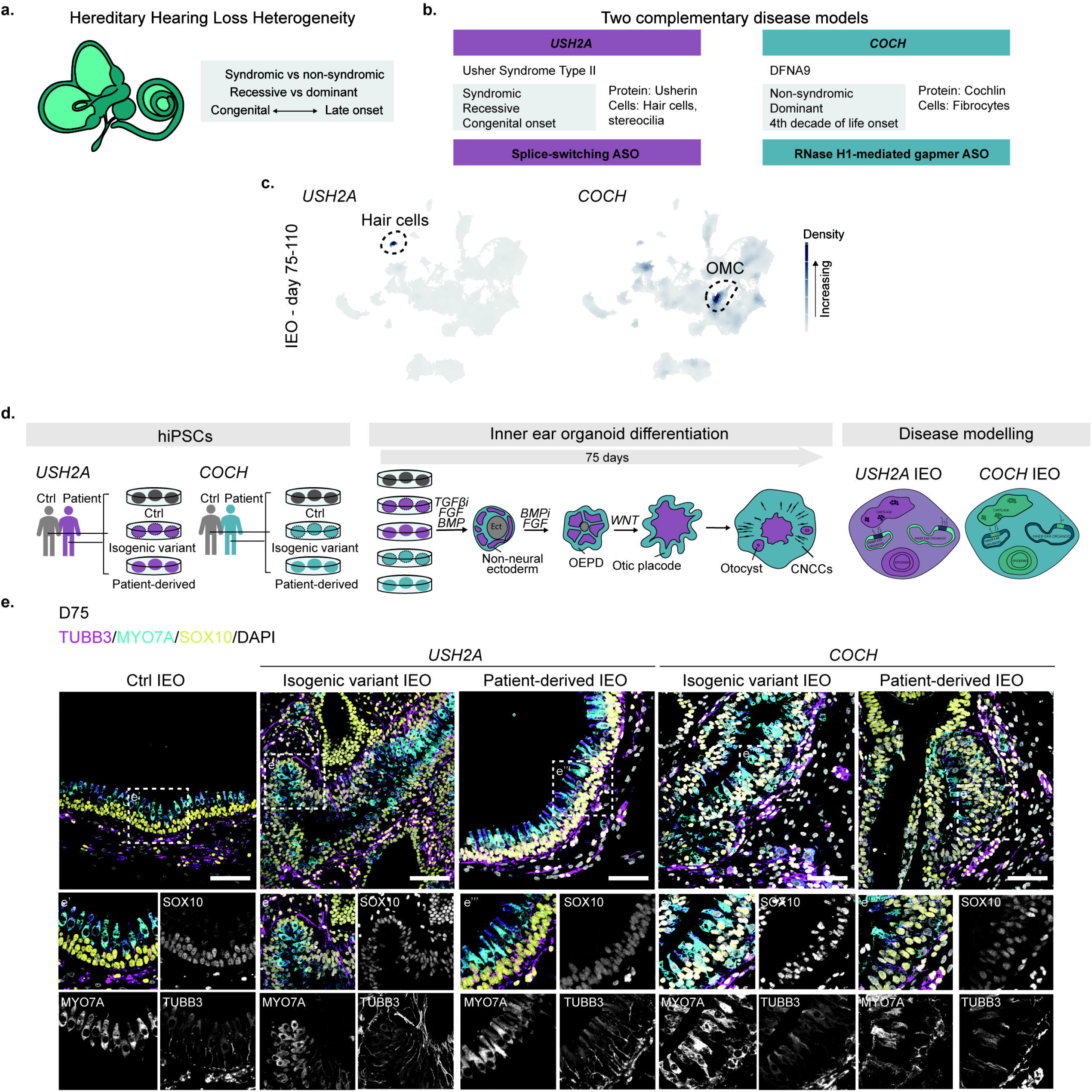
From Target Selection to Inner Ear Organoid (IEO) Modelling of Hereditary Inner Ear Diseases a. Visual representation of factors attributing to hereditary hearing loss heterogeneity (*2*). b. Overview of the presentation of auditory phenotypes in Usher Syndrome type II and DFNA9, as well as the localization of the disease-associated proteins usherin and cochlin, respectively. c. Feature plots showing gene expression of *USH2A* and *COCH* in single nuclei RNAseq data from day 75-110 human IEOs. d. Experimental set-up describing how healthy control, isogenic variant, and patient-derived induced pluripotent stem cells (iPSCs) were differentiated into IEOs to create *USH2A*-IEO and *COCH*-IEO for disease modelling. e. Representative panels showing immunofluorescence (IF) staining of day 75 IEOs derived from the control, isogenic variant and patient-derived iPSCs, containing otic vesicle-like structures (SOX10^+,^ yellow), hair cells (MYO7A^+,^ cyan) and neurons (TUBB3^+^, magenta). Scale bars: 50 µm.

To establish a system for personalized therapeutic testing, we generated IEOs from human induced pluripotent stem cells (hiPSCs) carrying pathogenic variants in *USH2A* and *COCH* (Figure 1d). Fibroblasts from two patients, one with Usher Syndrome Type II and one with DFNA9, carrying pathogenic variants in *USH2A* (c.7595-2144A>G, homozygous) or *COCH* (c.151C>T; p.Pro51Ser, heterozygous) were reprogrammed into iPSCs. In parallel, we made isogenic counterparts by introducing the corresponding *USH2A* homozygous or *COCH* heterozygous variant into a common healthy donor iPSC line, creating disease-control pairs on a shared genetic background. The respective genotypes for all iPSC lines were verified by Sanger sequencing (Figure S2).

Next, we differentiated all iPSC lines into IEOs. By day 75 of differentiation, all lines reproducibly formed inner ear-like structures characteristic of successful IEO differentiation (Figure 1e). The pathogenic variants did not impair organoid formation and the resulting organoids contained vesicles expressing SOX10, with both MYO7A^+^ hair cells and tubulin beta 3 class III (TUBB3^+^) neurons, confirming otic identity (Figure 1e).

This experimental design enabled us to successfully develop IEOs from iPSCs obtained from patients with genetically distinct hereditary inner ear diseases.

### *USH2A* variant IEOs recapitulate disease-associated *USH2A* pseudoexon inclusion

We next investigated whether IEOs retained the *USH2A* disease-causing variant within the relevant inner ear target cell populations (Figure 2). The pathogenic c.7595-2144A>G variant is located within an intronic region of *USH2A* and causes aberrant splicing leading to inclusion of a 152-bp pseudoexon 40 (PE40) into the mature transcript (Figure 2a, (44)). As pseudoexon inclusion is a defining molecular hallmark of this disease, we first assessed whether IEOs display the pathogenic splicing defect. Reverse Transcriptase (RT)-PCR analysis of day 75 IEOs detected the expected 1052-bp pseudoexon-containing transcript in both patient-derived and isogenic variant IEOs (Figure 2b).

**Figure 2.**
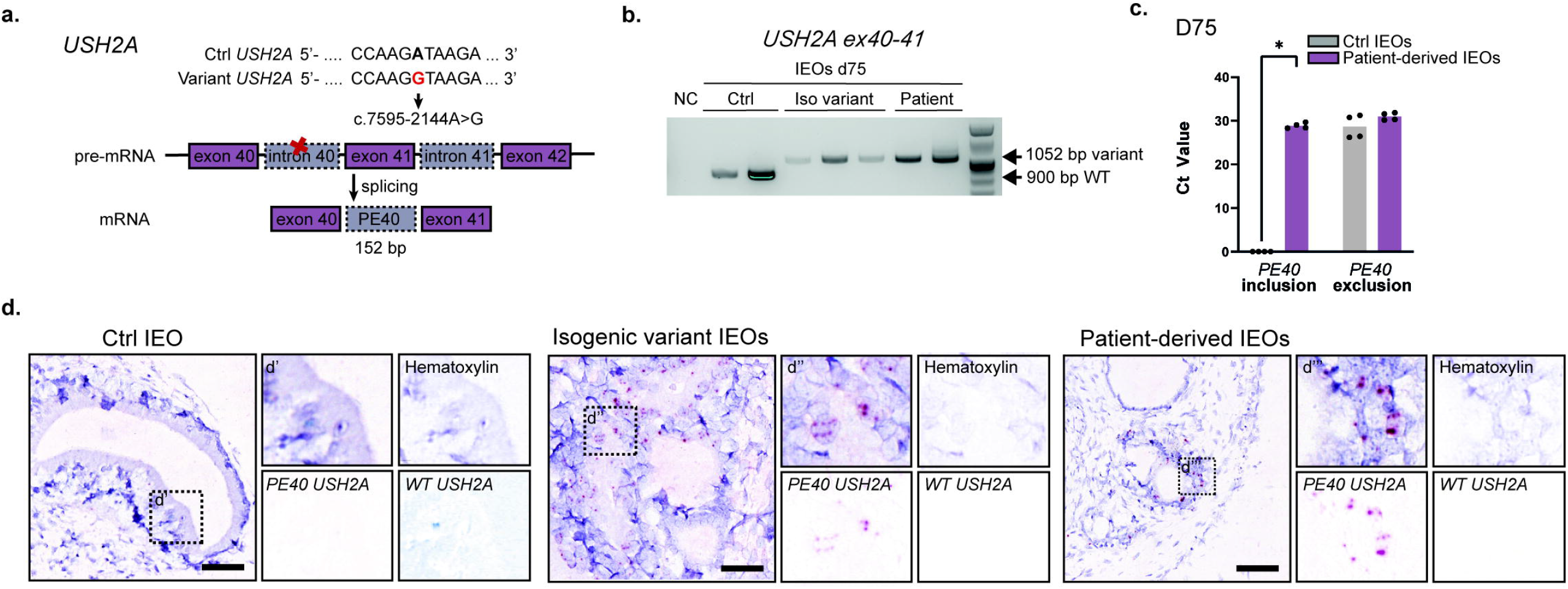
Patient-derived *USH2A*-variant IEOs Model Molecular Defects of *USH2A* pseudoexon inclusion a. Graphical explanation of the c.7595-2144A>G *USH2A* single nucleotide variant. b. Absence and presence of the pseudoexon-containing *USH2A* variant transcript in day 75 IEOs as shown by reverse transcriptase (RT)-PCR targeting *USH2A* ex40-41. c. qPCR of day 75 IEOs shows expression levels of pseudoexon-40 (PE40) inclusion and exclusion in control IEOs (*n* = 4) and patient-derived IEOs (*n* = 4). d. Images of in situ hybridization show the presence of wild type *USH2A* (blue) and pseudoexon-containing *USH2A* transcript (red) in day 75 IEOs. Scale bars: 50 µm.

Next, we quantified overall *USH2A* transcript abundance and observed differences between control, isogenic variant and patient-derived IEOs (Figure S3a). However, these differences appeared to correlate with variation in expression of the hair cell marker *MYO7A*, suggesting that they may reflect differences in hair cell abundance rather than changes in *USH2A* expression per se (Figure S3a). Consistent with hair cell differentiation, stereocilia bundle-like structures were observed by immunohistochemistry in both control and patient-derived IEOs at day 75 (Figure S3b). To further quantify pseudoexon inclusion, we used primer sets selective for PE40-containing transcripts and detected mutation-specific expression exclusively in disease-associated IEOs (Figure 2c).

To determine whether PE40 inclusion could be detected in the relevant inner ear cell populations, we performed *in situ* hybridization using probes specific for wild-type and PE40-containing *USH2A* transcripts. Transcript signal localized to the inner ear-like structures of the IEOs, consistent with the expected expression pattern of *USH2A* (Figure 2d). No PE40-containing transcript signal was detected in control IEOs, which exclusively expressed the wild-type transcript. In contrast, both isogenic variant and patient-derived IEOs showed PE40 containing transcript signal (Figure 2d). These results demonstrate that isogenic and patient-derived *USH2A*-variant IEOs reproduce the disease-associated transcript splicing defect characteristic of this early-onset inner ear disease.

### Modeling dominant hereditary hearing loss using *COCH* variant IEOs

To model a form of non-syndromic, dominantly inherited, late-onset hearing loss disorder, we generated IEOs from isogenic variant and patient-derived iPSC lines carrying the pathogenic *COCH* c.151C>T p.Pro51Ser variant associated with DFNA9 (Figure 3). The pathogenic *COCH* variant is heterozygous and located in exon 4 (Figure 3a), resulting in both wild-type and mutant transcripts being expressed.

**Figure 3.**
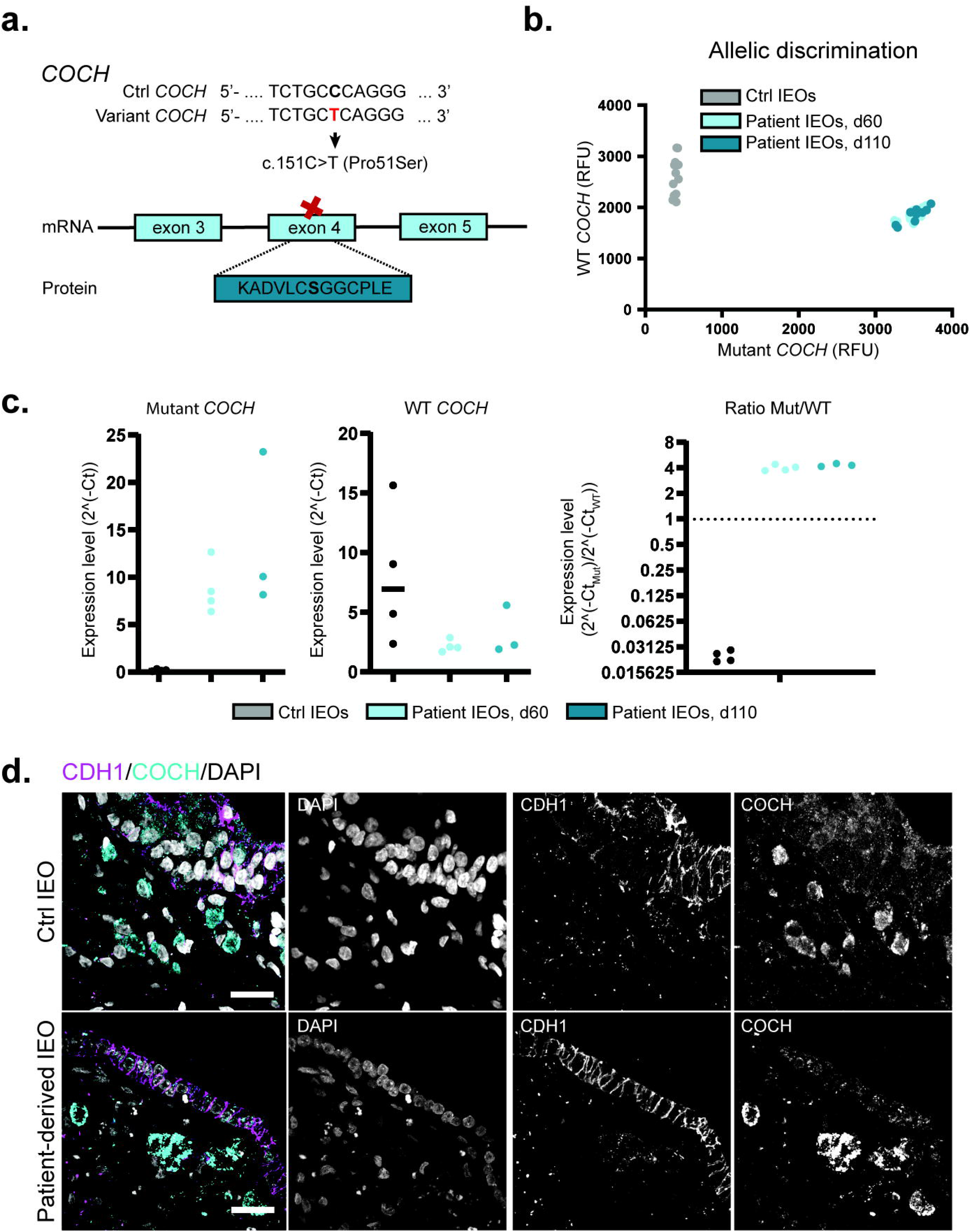
Patient-derived *COCH-*variant IEOs Model Molecular Defects. a. Graphical explanation of the *COCH* c.151C>T p.Pro51Ser variant. b. Allelic discrimination plot shows distinction between wild type and variant-containing allele of the *COCH* gene in day 75 control IEOs and patient IEOs (*n* = 4), as well as day 110 patient IEOs (*n* = 3), values indicate relative fluorescence units (RFU). c. Gene expression of the wild type and variant-containing alleles of the *COCH* gene in day 75 control IEOs and patient IEOs (*n* = 4), as well as day 110 patient IEOs (*n* = 3), values indicate relative gene expression levels from normalized RNA input in 2^(-Ct) The ratio is calculated as 2^(-Ct_MUT_)/ 2^(-Ct_WT_). d. Panels showing IF staining of day 75 control IEOs and patient-derived IEOs, containing cochlin^+^ cells (magenta) underlying the epithelial cells (CDH1^+,^ cyan). Scale bars: 20 µm.

To distinguish wild-type and mutant transcripts, we quantified allele-specific *COCH* expression using a previously established TaqMan assay (32). The TaqMan assays robustly discriminated between alleles (Figure 3b-c). Probe specificity was determined using synthetic wildtype and mutant gBlocks which excluded cross-reactivity between alleles (Figure S4a). Separate quantification of wild-type and mutant transcripts further confirmed preservation of the heterozygous allele expression within the IEOs (Figure 3c). Because DFNA9 develops progressively over time, we maintained IEOs in culture until both D75 and D110 to assess whether extended maturation would show features of DFNA9 pathology. We did not observe any differences between the two stages of differentiation. In addition, IEOs show robust overall *COCH* expression levels at both day 75 and 110, without significant temporal changes in transcript abundance (Figure S4b).

Given that DFNA9 pathophysiology is supposedly linked to the accumulation of cytotoxic cochlin aggregates in adult patient tissue, we next assessed cochlin localization by immunohistochemistry (Figure 3d). Cochlin is located in the mesenchymal-like compartment beneath the otic vesicle epithelium across all genotypes, consistent with periotic mesenchymal expression patterns (Figure 3d). However, we did not observe apparent cochlin aggregation in day 75 or day 110 IEOs, as described in post-mortem DFNA9 inner ear tissue (45). These findings likely reflect the fetal developmental state of the IEOs, which capture early molecular consequences of the pathogenic variant rather than end-stage pathology.

Together, these results demonstrate that patient-derived *COCH*-variant IEOs preserve the molecular features underlying DFNA9 and extend the IEO platform across distinct forms of hereditary inner ear diseases for therapeutic testing.

### ASOs delivery strategy and integrity assessment in IEOs

Having established these IEO models of hereditary inner ear disorders, we next investigated whether we could modify the observed disease-associated molecular defects using ASOs. Because the pathogenic mechanisms underlying *USH2A-*and *COCH-*associated inner ear pathologies differ fundamentally, we applied distinct ASO strategies for each mutation type.

For *USH2A* c.7595-2144A>G, we used a splice-switching ASO designed to redirect aberrant *USH2A* pre-mRNA splicing (Figure 4a, (46)). In contrast, for the *COCH* c.151C>T (p.Pro51Ser) variant, which exerts its dominant pathogenic effect via a non-haploinsufficiency disease mechanism, we used a gapmer ASO designed to recruit RNase H1 and selectively degrade mutation-specific transcripts (Figure 4b, (32)). The ASO sequences used here were identical to the ASOs reported in the original publications. Both ASOs are comprised of a fully phosphorothioated backbone, but the ribose groups contain 2’-O-methoxyethyl (MOE) modifications for the *USH2A* ASO and 2’-O-methyl (OMe) modifications for the *COCH* ASO.

**Figure 4.**
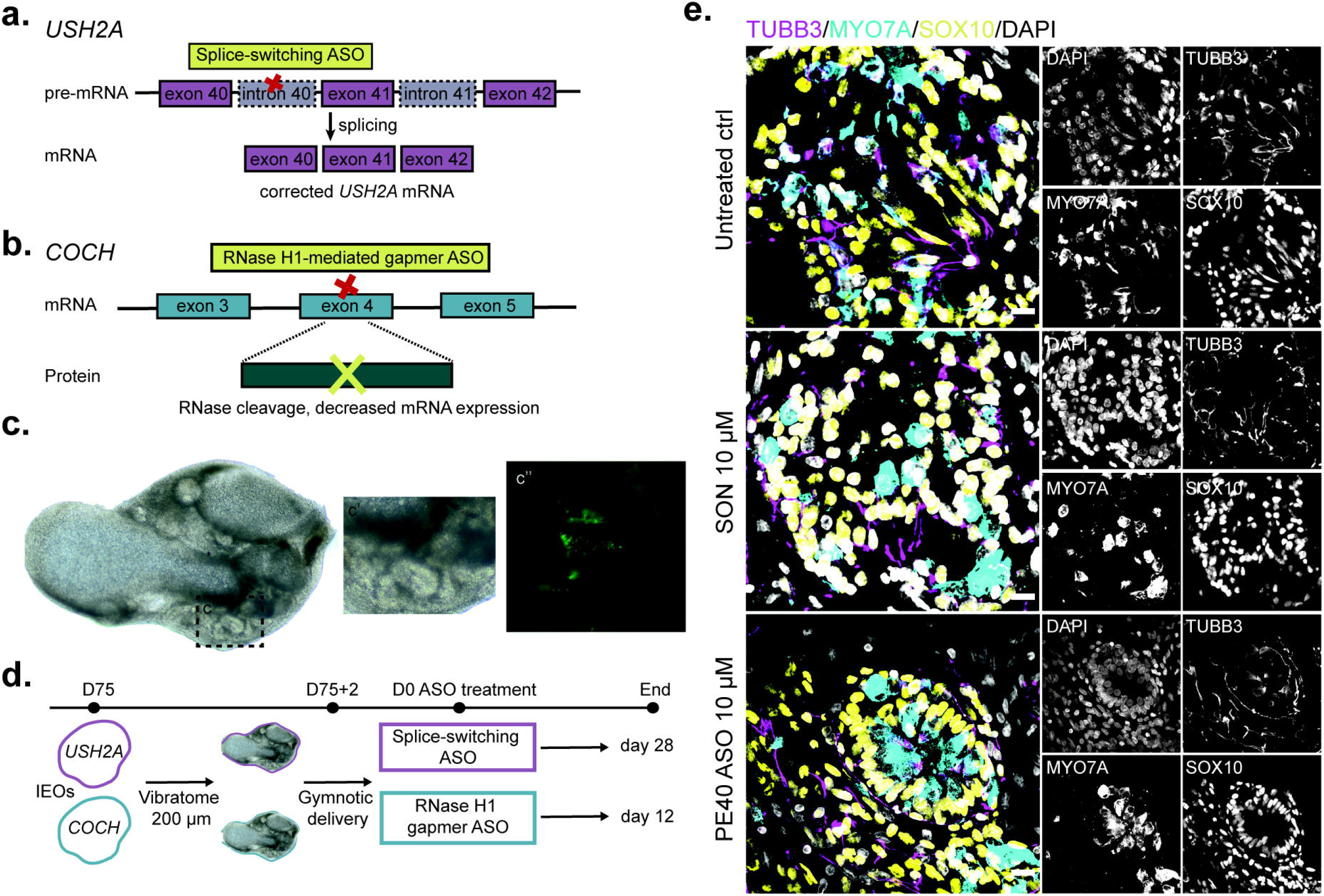
Delivery and Safety Assessment of Antisense Oligonucleotides in Inner Ear Organoids. a. Splice-switching ASO strategy for *USH2A* c.7595-2144A>G. b. Gapmer ASO strategy for *COCH* c.151C>T p.Pro51Ser. c. Example of a vibratome section showing (C’) otic vesicles and (C’’) stereocilia bundles (F-actin^+^, green). d. Experimental method of ASO delivery to IEOs. e. Representative panels showing immunofluorescence (IF) staining of *USH2A* patient-derived IEOs after 28 days treatment with 10 µM sense oligonucleotide (SON), 10 µM ASO and untreated, containing otic vesicle-like structures (SOX10^+,^ yellow), hair cells (MYO7A^+^, cyan) and neurons (TUBB3^+^, magenta). Scale bars: 20 µm.

To improve accessibility of ASOs to disease-relevant cell populations, we sectioned day 75 IEOs into 200 µm thick vibratome slices, thereby exposing target cells while preserving overall tissue architecture (Figure 4c). Additionally, this vibratome-based approach enabled us to identify the inner ear vesicles within the organoid and to confirm for instance hair cell presence with F-Actin staining prior to downstream analysis and reduced the impact of differentiation variability (Figure 4c’-c‘’). We subsequently maintained tissues in culture and directly administered naked ASOs to the medium (Figure 4d).

Next, we evaluated whether vibratome sectioning followed by high-dose ASO exposure altered organoid integrity. Immunohistochemical analysis of organoid slices with high-dose (10 µM) sense oligonucleotide (SON) or ASO showed preserved SOX10^+^ otic vesicle-like structures, MYO7A^+^ hair cells, and TUBB3^+^ neurons comparable to untreated controls (Figure 4e). Together, these findings indicate that there are no indications that the treatment conditions compromised overall tissue identity.

### ASO chemistry influences cellular uptake and intracellular localization in human IEOs

Because ASO efficacy depends on successful targeting and uptake by the appropriate cellular and intracellular compartments, we next investigated how different oligonucleotide chemistries distribute within IEOs. Although ASOs are increasingly explored as therapeutic strategies for sensory disorders, nothing is known about their uptake kinetics in the different human inner ear cell types. Understanding how distinct ASOs designs penetrate into disease-relevant target cells is essential for interpreting therapeutic efficacy and optimizing future inner ear-directed oligonucleotide therapies.

We first examined the distribution of the *USH2A* splice-switching ASO within patient-derived IEOs. Delivery of 10 µM Alexa Fluor 488-labeled PE40-ASO to vibratome slices resulted in progressive uptake throughout the tissue over the treatment period, ultimately resulting in broad distribution across the entire organoid slice (Figure 5a). Higher-resolution imaging confirmed uptake across multiple cellular populations within the IEOs (Figure 5b). Importantly, the labeled PE40-ASO reached the MYO7A^+^ hair cells, the primary disease-relevant target population for *USH2A*-associated disease (Figure 5b’-b’’). Within these cells, fluorescent signal localized to both the cytoplasm and nucleus, consistent with the intracellular access required for splice modulation.

**Figure 5.**
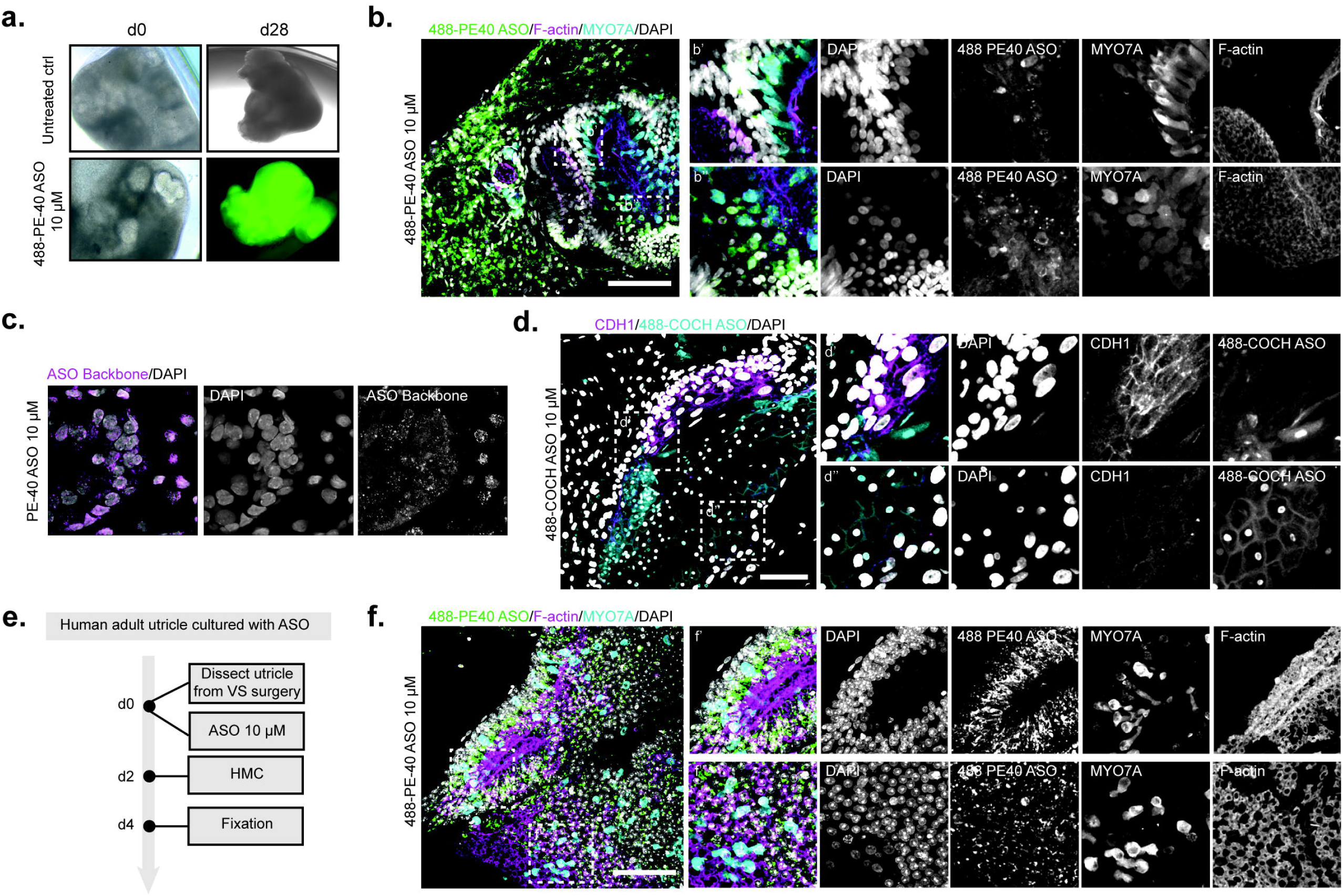
Distinct ASO Chemistries Show Differential Uptake in Inner Ear Organoids and Human Adult Utricle. a. Brightfield images of *USH2A* patient-derived IEOs after 28 days treatment with 10 µM Alexa Fluor (AF)488-conjugated splice-switching ASO. b. Representative panels showing IF staining of patient-derived IEOs after 28 days treatment with 10 µM 488-conjugated splice-switching ASO, containing the intrinsic signal of the AF488-PE40 ASO (green), hair cells (MYO7A+, cyan) and stereocilia bundles (F-actin^+^, magenta), with region 1 (C’) depicting a side view and region 2 (C”) depicting a top view. Scale bar: 50 µm. c. Representative panels showing IF staining of the localization of the ASO backbone (magenta) in patient-derived IEOs after 28 days treatment with 10 µM splice-switching ASO targeting pseudoexon-containing *USH2A* variant. Scale bar: 10 µm. d. Representative panels showing IF staining of patient-derived IEOs after 28 days treatment with 10 µM AF488-conjugated gapmer ASO, containing the intrinsic signal of the AF488-COCH ASO (cyan) and epithelial cells (CDH1^+^, magenta), with region 1 (E’) depicting a side view and region 2 (E”) depicting a top view. Scale bar: 50 µm. e. Experimental method of delivery and uptake of ASO therapy experiments with human adult utricle derived from vestibular schwannoma patients. f. Representative panels showing IF staining of human adult utricle after a 4-day treatment with 10 µM AF488-conjugated splice-switching ASO, showing the presence of the AF488 signal (green), hair cells (MYO7A^+^, cyan) and stereocilia bundles (F-actin^+^, magenta). Scale bars: 50 µm.

Since fluorophore conjugation can influence oligonucleotide uptake and intracellular trafficking, we next validated these observations using an antibody directed against the ASO’s phosphorothioate backbone in IEOs treated with unlabeled PE40-ASO (Figure 5c, Figure S5a). This independent approach confirmed successful cellular uptake and target cell access, with punctate intracellular staining detected within disease-relevant sensory regions. In addition, unlabeled PE40-ASO also localized to off-target cellular populations within the IEO tissue (Figure S5b). miRNAscope analysis targeting the PE40-ASO sequence reproduced these localization patterns (Figure S5c), collectively demonstrating efficient tissue penetration and intracellular delivery for the splice-switching oligonucleotide.

We next examined the *COCH* gapmer ASO, which differs from the splice-switching oligonucleotide in both chemistry and mechanism of action. Delivery of 10 µM AlexaFluor 488-labeled *COCH* gapmer ASO to patient-derived IEOs produced a distinct intracellular distribution pattern, characterized by a strong nuclear and plasma membrane-associated localization (Figure 5d). Notably, in contrast to the PE40-ASO, the fluorescent *COCH* gapmer showed limited uptake in CDH1+ cells of the sensory epithelium (Figure 4d, Figure S5d), suggesting that ASO chemistry and/or target cell identity strongly influence uptake behavior within IEOs.

### Uptake of ASOs is achieved in adult human vestibular tissue

To extend these findings beyond organoid tissue and evaluate the translational feasibility of inner ear antisense therapy, we next investigated ASO uptake in adult human vestibular tissue (Figure 5e-f). Adult human utricular tissue was obtained from patients undergoing translabyrinthine surgery for vestibular schwannomas.

To determine whether the *USH2A* splice-switching ASO could access disease-relevant sensory cells in mature tissue, adult utricles were treated with 10 µM fluorescently labeled PE40-ASO (Figure 5e). Immunohistochemical analysis demonstrated robust uptake within MYO7A^+^, F-Actin^+^ hair cells, indicating successful delivery of the oligonucleotide to the nucleus and cytoplasm of the relevant sensory compartment in adult human tissue (Figure 5f). Uptake patterns of PE40 ASO in adult utricular tissue were similar to uptake in IEOs. Together, these findings demonstrate that human IEOs provide a system to investigate ASO biodistribution and intracellular localization that may directly influence therapeutic performance.

### ASO treatment achieves target engagement in hereditary hearing loss IEOs

Having established ASO safety, tissue penetration, and cellular uptake within human IEOs, we next investigated whether ASO treatment could alter mutant transcripts in the IEO models (Figure 6). Because therapeutic efficacy depends not only on cellular uptake but also on sufficient intracellular exposure and transcript turnover, we first optimized treatment duration and dosing conditions for each genetic context.

**Figure 6.**
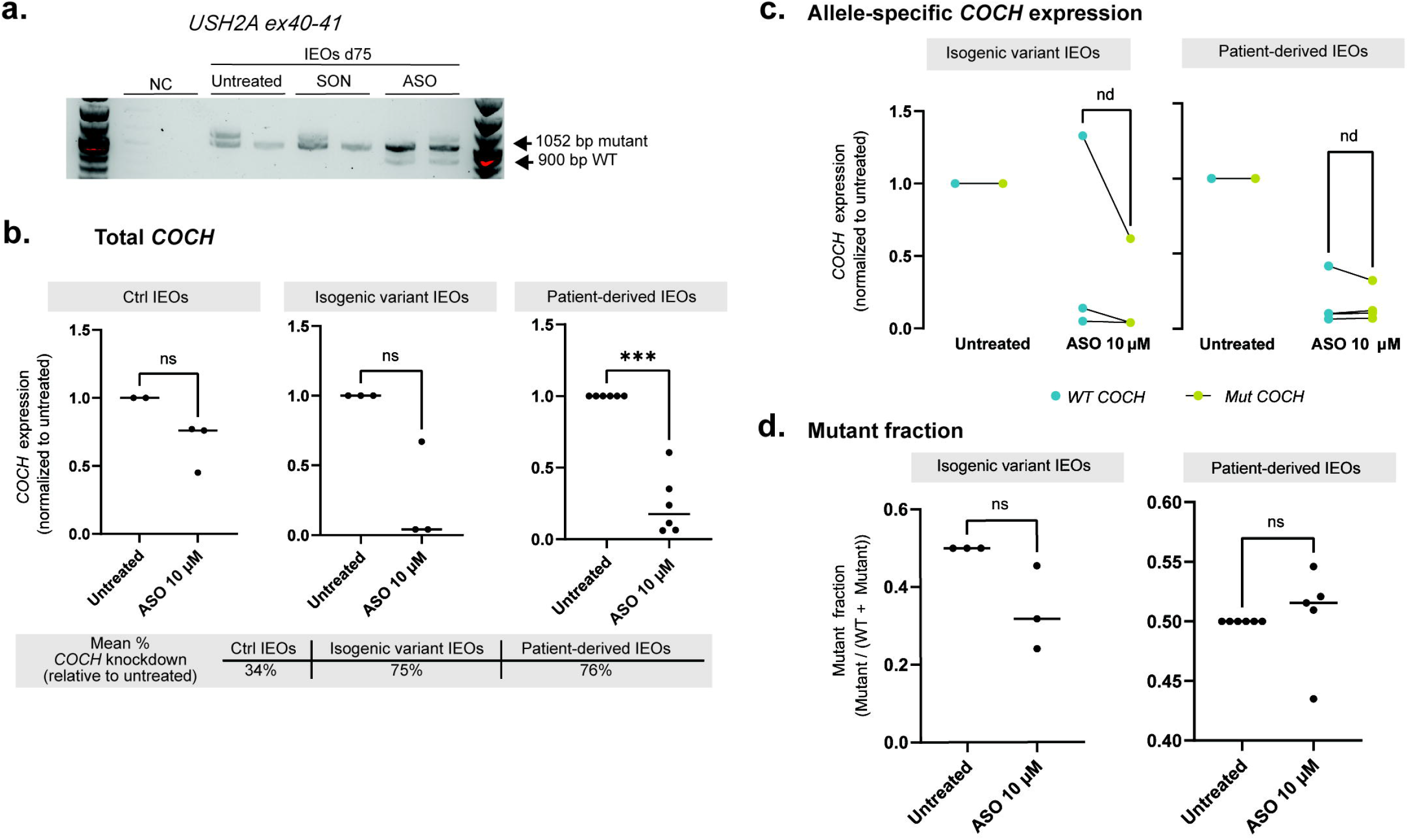
ASO Treatment Partially Restores Disease-Associated Molecular Defects Across Distinct Inner Ear Disease Models. a. Presence of wild type transcript in day 75 *USH2A*-patient-derived IEOs treated with 10 µM PE40-ASO, as shown by reverse transcriptase (RT)-PCR targeting *USH2A* ex40-41. NC indicates a negative control sample, SON indicates a nonbinding sense oligonucleotide. b. Total *COCH* expression decreases in patient-derived IEOs treated with 10 µM ASO, as shown by quantitative PCR (*n* = 6), expression levels are shown as 2^ (-dCT), normalized against untreated. Statistical test is Welch’s t-test (ns = not significant, ***p≤0.001). Mean percentage knockdown was calculated as (untreated – treated) * 100. c. Allele-specific wild-type (WT, blue) and mutant (Mut, green) *COCH* expression for isogenic variant IEOs (*n* = 3) and patient-derived IEOs (*n* = 6) in untreated and 10 µM ASO conditions, expression levels are shown as 2^ (-dCT), normalized to untreated samples. Statistical test is multiple paired t-tests (nd = non discovery). d. Mutant fraction was calculated for the wild-type and mutant *COCH* expression as mutant / (WT + Mutant). Statistical test is Welch’s t-test (ns = not significant, ***p≤0.001).

To define our initial treatment strategy, we performed experiments using a repeated dosing regimen over 12 days for both disease models, based on previous studies. In *COCH*-variant IEOs, this regimen produced the anticipated reduction in *COCH* transcript levels following gapmer ASO treatment. In contrast, 12 days of treatment did not sufficiently correct *USH2A* pre-mRNA splicing (Figure S6a). Because splice-switching ASOs act on newly synthesized pre-mRNA, and the turnover of *USH2A* pre-mRNA is thought to be long, we reasoned that prolonged ASO treatment may be required to obtain a detectable amount of splice-corrected transcripts. We therefore extended treatment duration to 28 days for subsequent *USH2A* experiments.

For *USH2A*, we applied the splice-switching ASO designed to prevent inclusion of pseudoexon 40 and restore formation of the wild-type transcript. Using the previously introduced RT-PCR assay, we detected partial recovery of the 900-bp wild type transcript in patient-derived IEOs treated with 10 µM ASO, whereas SON-treated IEOs retained the aberrant pseudoexon-containing transcript (Figure 6a). These findings demonstrate partial correction of the pathogenic splicing defect in the IEO model.

For *COCH*, we applied the gapmer ASO strategy consistent with the dominant-negative nature of the mutation (Figure 6b-e, Table 1). In this setting, the therapeutic goal was preferential depletion of the transcripts containing the pathogenic variant while preserving expression of the wild-type allele. As expected, healthy control IEOs showed no significant reduction following treatment with 10 µM ASO (Figure 6b), although when quantified a small reduction of *COCH* expression was observed (34%). Isogenic variant IEOs showed a stronger *COCH* reduction upon 10 µM ASO treatment (75%), but this reduction varied between the treated samples, leading to an overall non-significant difference when compared to untreated controls. This could be due to the small sample size as well as the inherent variability in initial *COCH* expression between samples. In patient-derived IEOs, the 10 µM ASO conditions demonstrated a significant reduced expression of total *COCH*. In isogenic variant and patient-derived IEOs treated with a scrambled ASO (SCR) a reduction of *COCH* expression was also observed (Figure S6b-d).

**Table 1.** Antisense oligonucleotide sequences used in this study.

| Name | Sequence | Chemistry | Company | Reference |
| --- | --- | --- | --- | --- |
| USH2A_PE40_ASO_Fluo | C*C*C*-T*T*A*-A*A*A*-G*C*C*-A*G*C*-A*T*A*-C*A | MOE | Eurogentec | n/a |
| USH2A_PE40_ASO | C*C*C*-T*T*A*-A*A*A*-G*C*C*-A*G*C*-A*T*A*-C*A | MOE | Eurogentec | (46) |
| USH2A_SON | T*G*T*-A*T*G*-C*T*G*-G*C*T*-T*T*T*-A*A*G*-G*G | MOE | Eurogentec | (46) |
| COCH_E1_Fluo | mA*mG*mC* mC*C*C* C*T*G* A*G*C* A*G*A*<br>G*mG*mA* mC*mA | DNA/OME | SynOligo | n/a |
| COCH_E1 | mA*mG*mC* mC*C*C* C*T*G* A*G*C* A*G*A*<br>G*mG*mA* mC*mA | DNA/OME | Eurogentec | (32) |
| COCH_SCR | mG*mC*mU* mA*T*C* G*A*T* T*A*C* A*C*T*<br>A*mU*mC* mG*mA | DNA/OME | Eurogentec | (32) |

To further define allele specificity, we next performed allele-specific TaqMan assays to separately quantify wild-type and mutant *COCH* transcripts. ASO exposure reduced both wild-type and mutant transcript levels, in the majority of the treated samples, indicating diminished allele specificity at these exposure levels (Figure 6c). As a result, the mutant fraction in both isogenic variant and patient-derived IEOs did not change after the addition of ASO 10 µM (Figure 6d).

Together, these findings demonstrate that ASO therapy partially correct disease-associated molecular defects in both *USH2A*- and *COCH*-variant IEOs, establishing the therapeutic generalizability of this platform across mechanistically distinct forms of hereditary inner ear diseases.

## Discussion

Progress toward precision therapies for hereditary inner ear disorders has been constrained by limited access to human inner ear tissue, the limited throughput and translatability of existing animal models, and the absence of scalable human in vitro systems that capture differentiated inner ear target cells and tissue architecture. In this study, we establish human IEOs as a platform for both mutation-specific disease modeling and preclinical evaluation of ASOs. We generated disease models representing genetically and clinically distinct forms of hearing loss, with syndromic and non-syndromic, and early- and late-onset disease contexts, and therapeutic approaches based on either splice-switching ASOs or RNase H1-mediated transcript degradation using gapmer ASOs. Disease-specific IEOs enabled the evaluation of ASO activity in different target cell populations, including hair cells and otic mesenchyme cells, while also assessing the uptake of distinct ASO chemistries.

To model hereditary inner ear disease, we used two complementary strategies for generating hereditary inner ear disease models: the introduction of pathogenic variants into established iPSC lines and the use of patient-derived iPSCs. Both approaches successfully generated disease-relevant human models and offer distinct advantages for therapeutic development. Isogenic disease-control models enable controlled investigation of variant-specific effects, while patient-derived models capture the broader genetic context in which disease occurs. The ability to combine these approaches within the same experimental framework provides a flexible platform for studying various hereditary inner ear disease models and evaluate personalized therapeutic interventions.

Beyond disease modeling, a major challenge in inner ear therapeutic development is determining whether therapeutic molecules can effectively reach disease-relevant cell populations within the largely inaccessible human inner ear. Human data on ASO biodistribution within inner ear tissue remain scarce. Here, we show that IEOs provide an opportunity to address this question preclinically by enabling visualization of ASO uptake in disease-relevant cell populations within three-dimensional tissues. To preserve tissue architecture and spatial information, we prioritized a vibratome-based workflow over dissociation-based approaches. Although this approach has a lower throughput rate than dissociation-based (28) or transfection-assisted (30) methods and might be less suited for large-scale ASO screening, it provides insights into ASO distribution within complex human tissues. In addition, the vibratome-based approach improved ASO uptake by exposing target cell populations at the surface, and this delivery method somewhat resembles a potential future delivery strategy for the human inner ear in the future. As RNA-based therapeutics continue to advance, the evaluation of tissue delivery and target cell uptake will remain an important area of research.

Using this approach, we evaluated ASO uptake in both human IEOs and adult human vestibular tissue and found that ASO chemistry influenced tissue penetration, cellular uptake and intracellular localization. We found that the splice-switching *USH2A* ASO efficiently reached hair cells and localized to both the cytoplasm and nucleus, consistent with the intracellular compartment required for modulation of pre-mRNA splicing and previous *USH1C* mouse studies (33). In contrast, the gapmer *COCH* ASO displayed a distinct uptake pattern with prominent nuclear and membrane-associated localization and more restricted uptake in non-epithelial populations, indicating that the ASO chemistry and mechanism of action may influence cellular uptake. To the best of our knowledge, studies examining biodistribution of gapmer ASOs in a human inner ear context have not previously been reported. Jang et al (2025 (47)) did show that gapmer ASOs directed against a mutant transcript of *KCNQ4*, highly expressed in outer hair cells (OHCs), was able to improve auditory function upon the intracochlear injection in pups of their DFNA2 mouse model. While difference between species and developmental stages could explain this, *KCNQ4* is also expressed in neuronal and lateral wall cells of adult mice. Because ASO biodistribution was not investigated in Jang et al, it is premature to conclude the effect they observed results from ASO activity in OHCs. As such, our work here could be a starting point for future investigations into the determinants of ASO delivery and activity in specific cell types of the inner ear.

The ability to evaluate cellular uptake and molecular efficacy within the same human tissue model represents a major advantage of the IEO platform. For the splice-switching *USH2A* ASO (46), molecular rescue could also be shown in the IEO through restoration of correctly spliced transcripts, providing a direct and internally controlled measure of therapeutic activity. In contrast, evaluation of RNase H1-mediated transcript degradation with gapmer ASOs required allele-specific strategies. Using allele-specific TaqMan assays, we observed reduced expression of mutant-containing transcripts, demonstrating target engagement by the *COCH* ASO in the IEOs, in line with previous studies (32). With the higher cumulative dosing used in this study, however, reduction of wild-type transcript expression was also observed, highlighting the importance of optimizing ASO chemistry, dosing and exposure to optimize allele-specificity. More broadly, splice-switching and gapmer ASOs present distinct analytical challenges. Whereas splice-switching ASOs inherently contain an internal molecular control through restoration of normal transcript products, gapmer-mediated degradation strategies rely heavily on external quantitative controls and highly sensitive allele-specific assays. For gapmers targeting *COCH*, variability in the proportion of *COCH*-expressing cells between IEOs could influence measured transcript levels and affect the interpretation of ASO-mediated knockdown. Future implementation of digital droplet PCR, or identification of stable, target cell-specific transcripts for the normalization of ASO effects, can improve sensitivity for detecting subtle allele-specific effects and therapeutic specificity (48). Overall, efforts to develop more advanced techniques to evaluate gapmer ASO effecicacy in IEOs are needed (49).

Importantly, our findings suggest that IEOs are well suited for evaluating therapeutic interventions in a patient-specific molecular context. By enabling assessment of target engagement, transcript correction, and cellular uptake within human inner ear cell populations, IEOs fill a gap between conventional cell culture systems and in vivo animal models. Similar to developments in the retinal field, where organoid systems are increasingly recognized as translational platforms that complement animal models and provide reproducible human-specific insights into therapeutic efficacy (50–52), human IEOs have the potential to occupy a comparable position within inner ear therapeutic development. Rather than replacing existing preclinical models, organoids may serve as an intermediate step that bridges genetic diagnosis, mechanistic investigation, and therapeutic testing in human tissue.

Despite these advances, several limitations of the current study should be considered. While the organoids successfully captured disease-relevant molecular defects and enabled assessment of therapeutic responses, the present study focused primarily on molecular endpoints and did not evaluate functional loss or rescue. Future studies incorporating electrophysiological and mechanosensory assays will be necessary to determine whether molecular correction translates into restoration of cellular function. In that case, cochlear IEOs are more suited to answer these questions (53) than the vestibular IEOs used in this study, as cochlear organoids have the possibility for functional assessment of hearing. In addition, batch-to-batch variability remains an inherent challenge of human IEO differentiation, and variability in the abundance and spatial distribution of target cell populations complicates quantitative morphological analyses. Current organoid maturation states may also limit modelling of age-dependent or late-onset disease phenotypes and frequency-specific hearing loss. This is particularly relevant for disorders such as DFNA9, in which pathogenic mechanisms, including cytotoxic cochlin aggregation, develop progressively over decades and may not be fully captured in fetal-like organoid models. In addition, the lack of tonotopic organization in current IEO systems limits their ability to model frequency-specific patterns of hearing loss. Continued optimization of differentiation protocols, maturation strategies, and quantitative analytical methods will further enhance the use of IEOs for therapeutic development. Finally, important biological questions remain in translational research that cannot be answered with IEOs such as the relationship between pathogenic variants and disease progression, the optimal timing of therapeutic intervention, and the duration of potential therapeutic windows in different forms of hereditary hearing loss.

ASOs are particularly well-suited therapeutic candidates for hereditary inner ear disease caused by pathogenic variants with a dominant negative or toxic gain of function effect. These disease mechanisms are often challenging to address with gene replacement or other DNA-based approaches (54). In addition, ASOs do not require viral vectors, thereby avoiding challenges associated with vector packaging limits, manufacturing and long-term side effects. This may be particularly beneficial for disorders caused defects in by large genes or in genes expressed across multiple inner ear cell populations, where broad distribution of a therapeutic molecule may be required. Consistent with this, our studies show uptake of ASOs in disease-relevant cell types within human inner ear tissue.

However, substantial translational hurdles also remain before ASO-based therapies for hereditary hearing loss can be implemented in clinical practice. Current organoid systems do not capture pharmacokinetic and pharmacodynamic parameters that will be essential for defining dosing intervals, long-term tissue persistence, and therapeutic safety. The relationship between the ASO concentration and effective exposure levels within the human inner ear also remains unclear. In addition, optimal timing of intervention may differ substantially between congenital and adult-onset disorders. Repeated ASO administration may be required in patients, raising important considerations regarding treatment burden, long-term adherence, and cost-effectiveness relative to existing standards of care such as cochlear implantation. Continued development of delivery approaches, including local slow-release systems, may help address some of these challenges. Finally, implementation of individualized ASO therapeutics will likely require continued development of regulatory pathways capable of supporting mutation-specific or N-of-1 therapeutic development (55,56).

In conclusion, our study demonstrates the potential of human IEOs as a platform for therapeutic development in hereditary inner ear disease. By combining patient-derived and isogenic disease models with ASO therapies, we established a workflow that enables the study of hereditary inner ear diseases and molecular rescue in a human inner ear context. Importantly, this approach was applicable across genetically and clinically distinct diseases, highlighting the versatility of organoid-based platforms. As organoid maturation, availability of functional readouts, and delivery strategies continue to improve, human IEOs will become increasingly important models to bridge the gap between genetic diagnosis and individualized treatment. Together, these findings provide a foundation for the development and preclinical evaluation of precision therapies for a broad range of hereditary inner ear diseases.

## Declarations

## Ethics approval and consent to participate

Patient fibroblasts were retrieved and reprogrammed into iPSCs under approval of Local Ethics Committee Radboud UMC (Centrale Commisie Mensgebonden Onderzoek, CMO-light, protocol number 2015-1543). The collection of human vestibular samples during translabyrinthine vestibular schwannoma surgery was approved by the Medical Research Ethics Committee of Leiden University Medical Center (protocol registration number BB23.001) and obtained after written informed consent of the donor.

## Consent for publication

n/a

## Availability of data and materials

Scripts used for the human inner ear dataset are available at https://github.com/OtoBiologyLeiden/Human-Fetal-and-Adult-Inner-Ear-Atlas. The raw single-nucleus RNA-seq data have been deposited at the Gene Expression Omnibus (GEO: GSE213796) by the authors of the original article (19). All other data supporting the findings of this study are available within the paper and its Supplementary Information. For further access to the data please contact the corresponding author.

## Authors’ contributions

- Conceptualization: WvdV, HK, EvW, EdV, HL
- Methodology: EF, WvdB, WvdV, EdV, HL
- Software: EF, WvdB
- Validation: EF, WvdB, EvW, WvdV, EdV, HL
- Formal analysis: EF, WvdB, EdV
- Investigation: EF, WvdB, AL, SU
- Resources: EF, WdvB, HK, EvW, WvdV, EdV, HL
- Data curation: EF, WvdB
- Writing – original draft: EF, WvdB, WvdV, HL
- Writing – review & editing: all authors
- Visualization: EF, WvdB
- Supervision: WvdB, HL
- Project administration: EF, WvdB, PPvB, WvdV, HL
- Funding acquisition: WvdV, HL

## Acknowledgments

We thank M. Bax for providing the vibratome, B. van der Vaart and C. Freund (LUMC hiPSC Centre) for creating the isogenic variant iPSC lines, the LUMC Light Microscopy Facility for microscopy assistance, and LUMC ENT surgeons E.F. Hensen and J.C. Jansen for providing human vestibular tissue. HK, EvW and EdV are members of the European Reference Network for rare and/or complex craniofacial anomalies and ear, nose and throat (ENT) disorders, ERN CRANIO.

## Supplementary Figures

**Figure S1.**
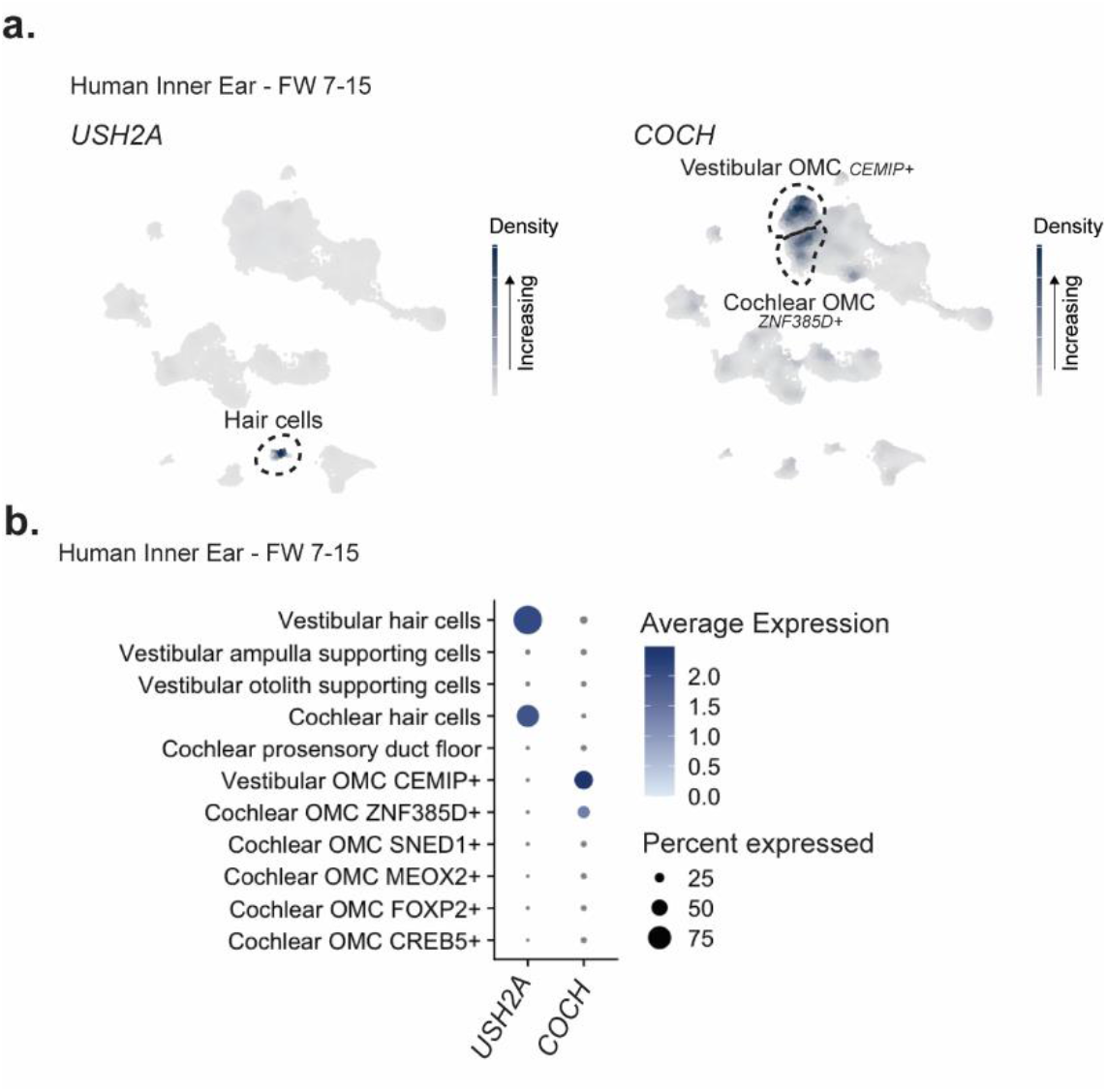
Expression of *USH2A* and *COCH* in human fetal inner ear. a. Feature plots showing gene expression of *USH2A* and *COCH* in single-nuclei RNAseq data from fetal week 7-15, retrieved from Human Inner Ear Development single cell RNA Atlas (HIEDRA, (41)). b. Dot plot showing *USH2A* and *COCH* gene expression levels in relevant inner ear cell types, indicating the average relative expression among expressing cells (dot color) and the percentage of cells expressing the gene within each annotated cell type (dot size).

**Figure S2.**
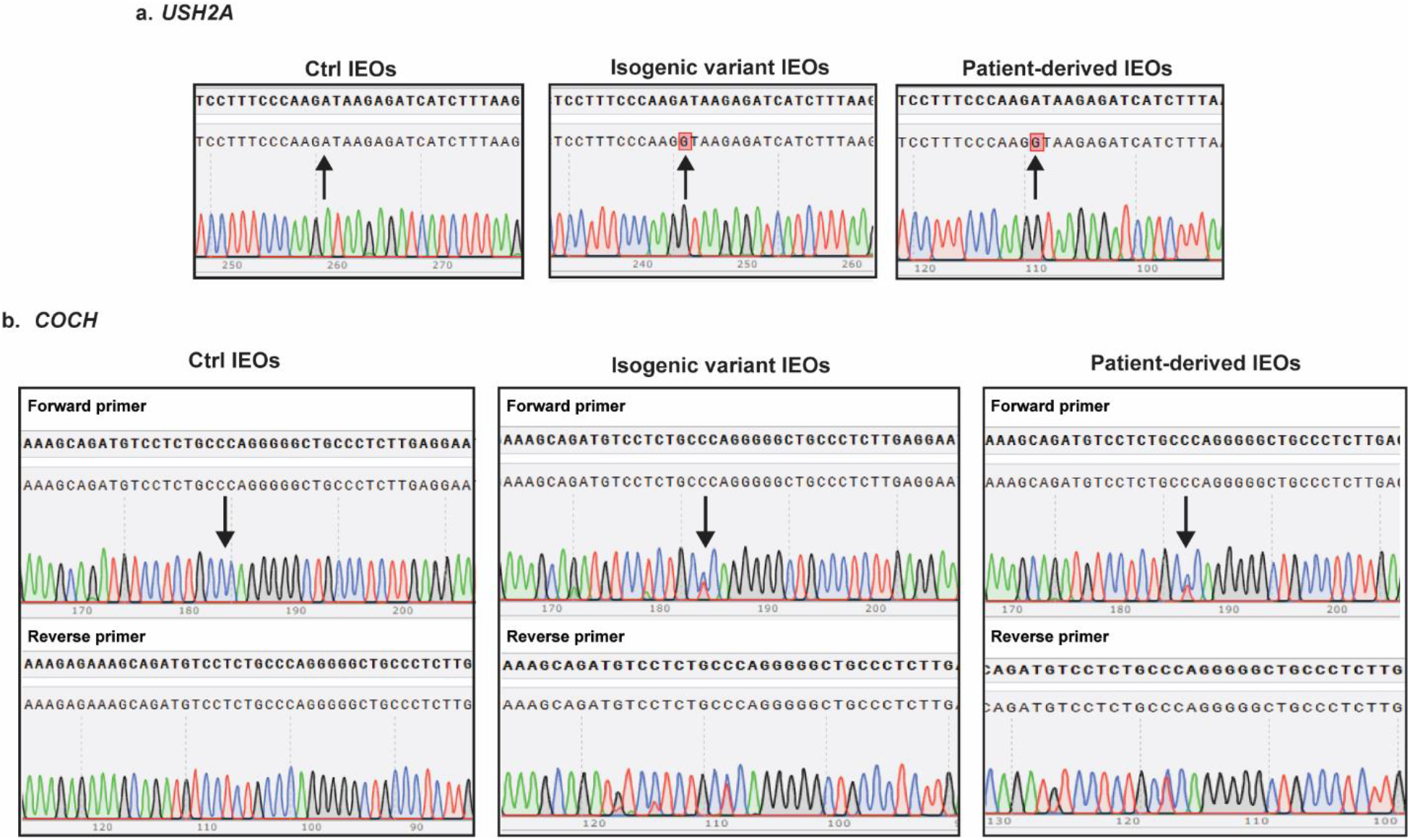
Confirmation of genotypes with Sanger Sequencing A. *USH2A* (c.7595-2144A>G, homozygous) in control, isogenic variant and *USH2A-*patient derived IEOs. Arrows indicate the site of the variant. B. *COCH* (c.151C>T; p.Pro51Ser, heterozygous) in control, isogenic variant and *COCH*- patient derived IEOs. Arrows indicate the site of the variant.

**Figure S3.**
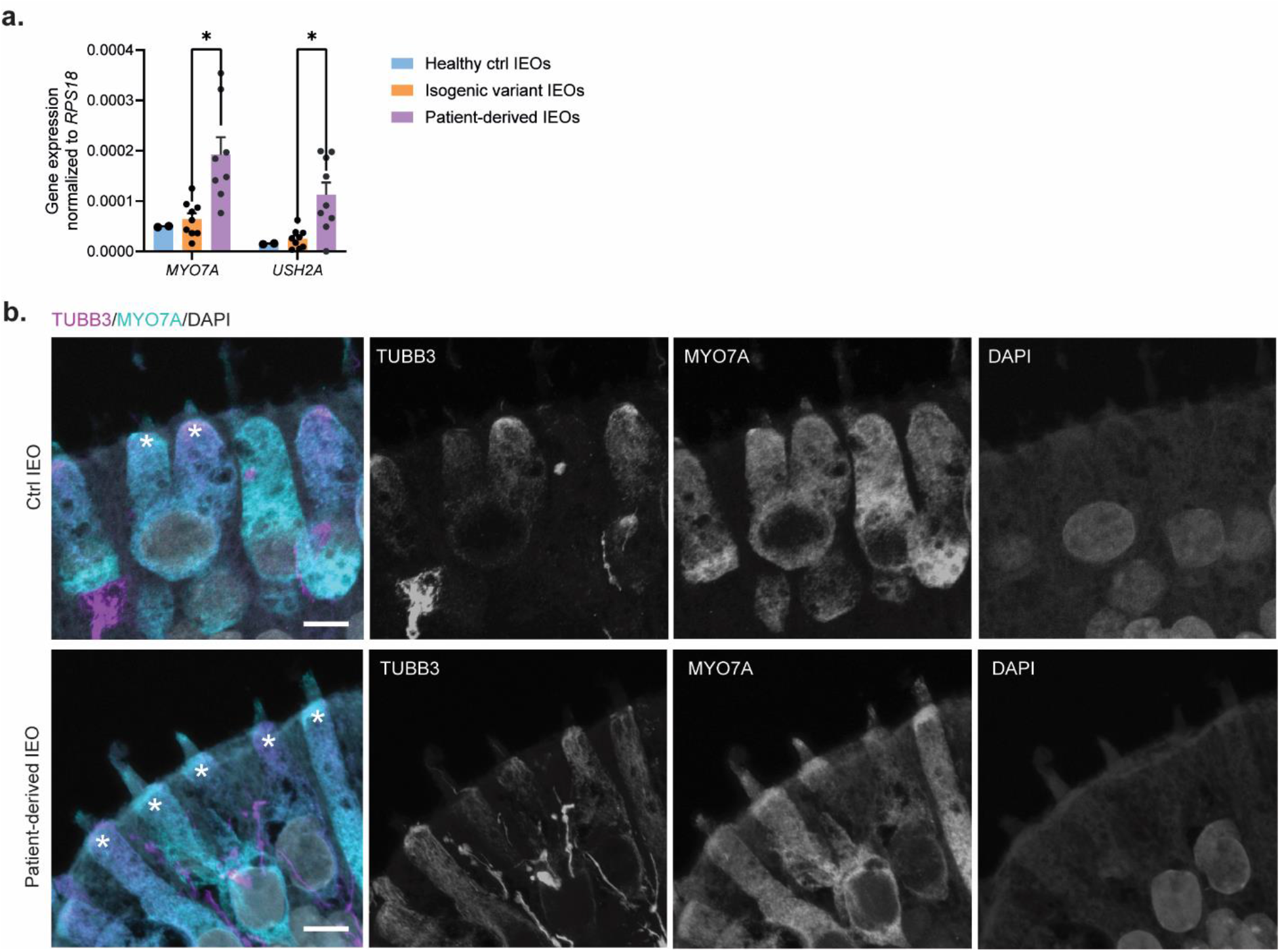
Additional information *USH2A* a. Gene expression of *MYO7A* and *USH2A* in day 75 control (*n* = 2), isogenic variant (*n* = 9) and *USH2A* patient-derived (*n* = 9) inner ear organoids (IEOs), normalized to *RPS18* expression. Statistical test via multiple unpaired t-tests with Welch correction (*p≤0.001). b. Representative panels showing immunofluorescence (IF) staining of day 75 IEOs derived from the control and *USH2A* patient-derived iPSCs, containing hair cells (MYO7A^+^, red) and neurons (TUBB3^+^, yellow). * indicate the base of the stereocilia bundle. Scale bars: 5 µm.

**Figure S4.**
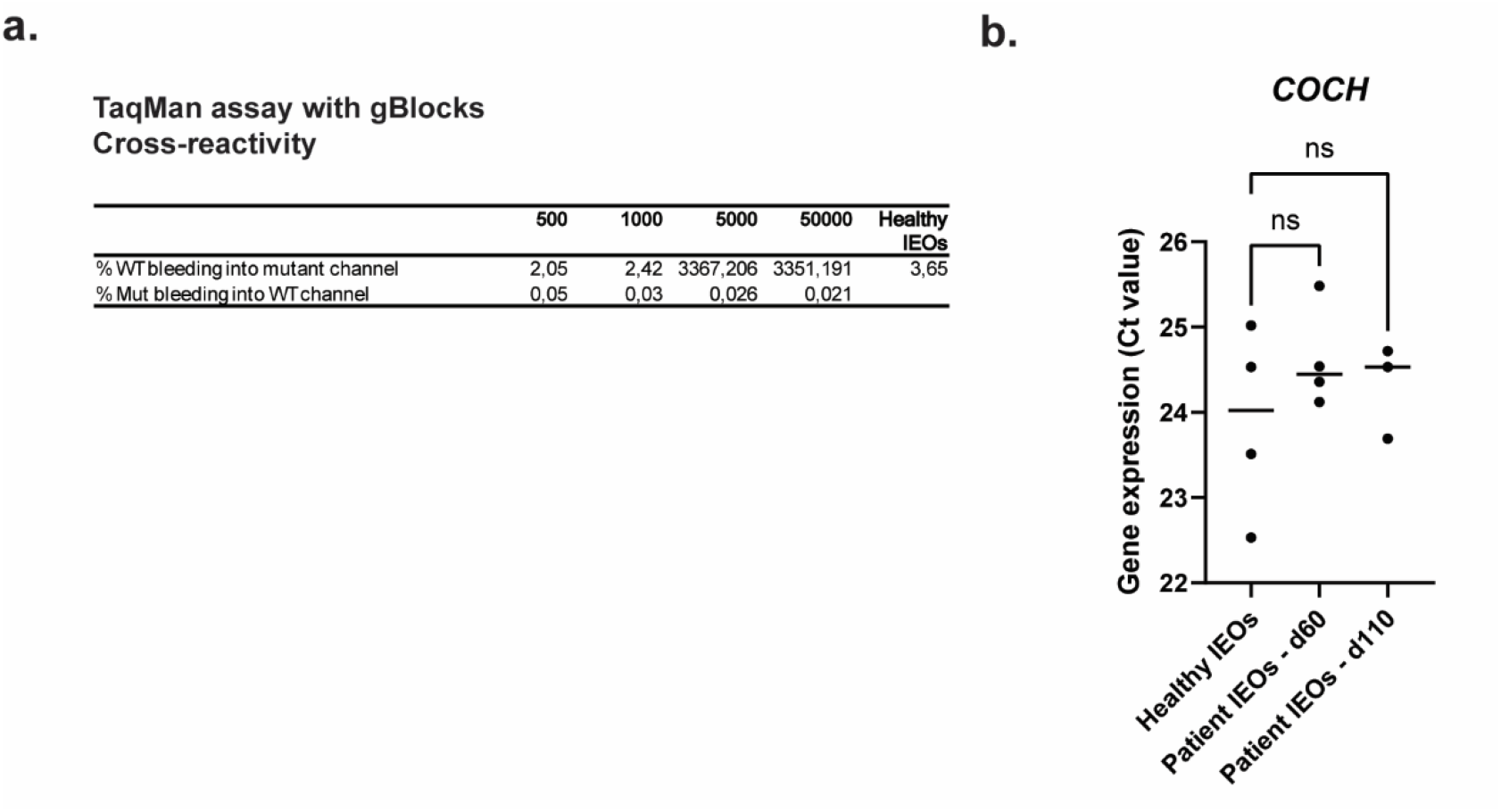
Additional information for *COCH* a. Cross-reactivity coefficients calculated from TaqMan assay wild type (WT) and mutant *COCH* c.151C>T expression using gBlocks of wild type and mutant *COCH* in 500-50000 copies/µl. Cross-reactivity = 2^(-A) / 2^(-B) * 100, in which A = ‘Ct value_gBlock WT – Mut channel’ and B = ‘Ct value_gBlock WT – WT channel’. b. Gene expression (Ct value) of the *COCH* gene upon normalized RNA input in day 75 control (*n* = 4), patient IEOs day 60 (*n* = 4) and patient IEOs day 110 (*n* = 3). Statistical test via ordinary one-way ANOVA (ns = non-significant).

**Figure S5.**
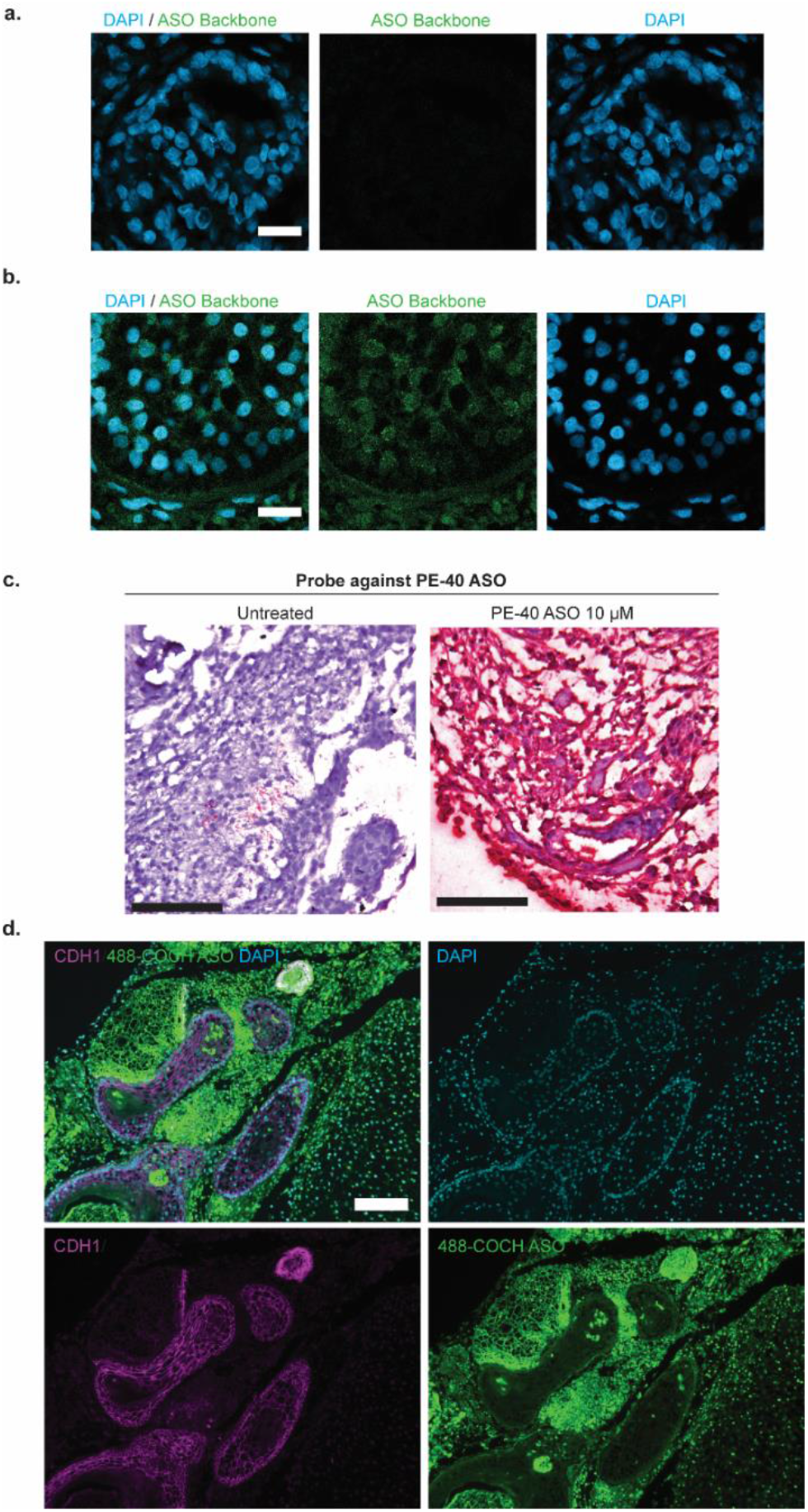
Additional ASO uptake experiments. a. Representative panels showing negative control immunofluorescence staining of the localization of the ASO backbone (green) in off-target tissues in patient-derived IEOs after 28 days treatment with 10 µM splice-switching ASO targeting pseudoexon-containing *USH2A* variant. Scale bar: 20 µm. b. Representative panels showing immunofluorescence staining of the localization of the ASO backbone (green) in off-target tissues in patient-derived IEOs after 28 days treatment with 10 µM splice-switching ASO targeting pseudoexon-containing *USH2A* variant. Scale bar: 20 µm. c. Probe against PE40 *USH2A* ASO shows localization of PE40-ASO throughout the IEO treated with 10 µM splice-switching ASO. Scale bar: 100 µm. d. Representative panels showing IF staining of the localization of the AF488-labeled gapmer ASO targeting COCH, showing that the labeled ASO (green) does not localize in epithelial cells (CDH1^+^, magenta). Scale bar: 100 µm.

**Figure S6.**
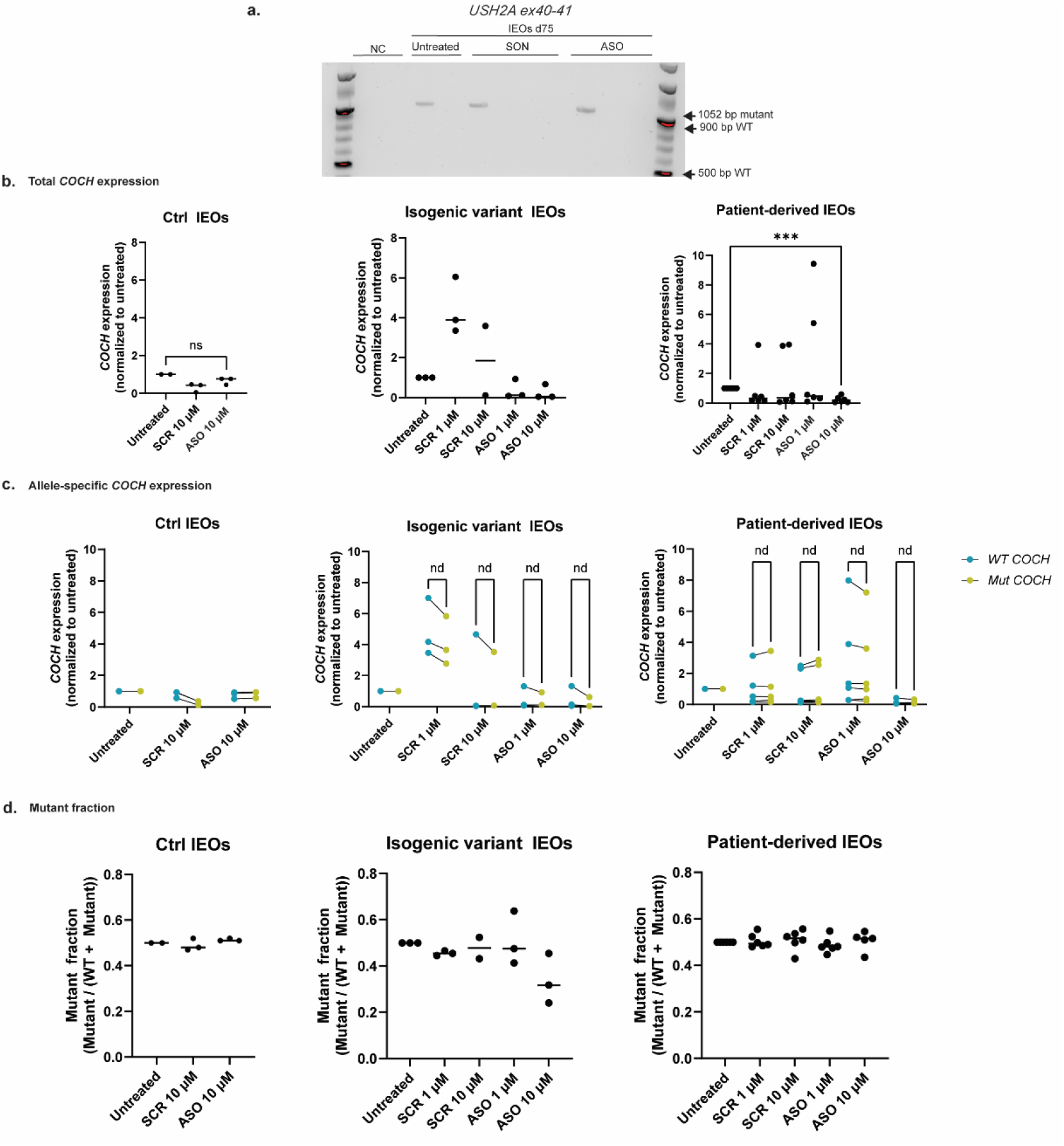
Total *COCH* and allele-specific *COCH* expression in ctrl IEOs, isogenic variant IEOs and patient-derived IEOs. a. Absence of wild type transcript in day 75 *USH2A*-patient-derived IEOs treated with PE40-ASO for 12 days, as shown by reverse transcriptase (RT)-PCR targeting *USH2A* ex40-41. NC indicates a negative control sample, SON indicates a nonbinding sense oligonucleotide. b. Total *COCH* expression in control IEOS (*n* = 3), isogenic variant IEOs (*n* = 3) and patient-derived IEOs (*n* = 6) as shown by quantitative, normalized to the untreated sample. Statistical test via Welch’s t-test (ns = not significant, ***p≤0.001). c. Allele-specific wild-type (WT, blue) and mutant (Mut, green) *COCH* expression for control IEOs (*n* = 3), isogenic variant IEOs (*n* = 3) and patient-derived IEOs (*n* = 6) in untreated and ASO conditions, normalized to untreated samples. Statistical test via multiple paired t-tests (nd = non discovery). d. Mutant fraction was calculated for the wild-type and mutant *COCH* expression as mutant / (WT + Mutant). Statistical test via Welch’s t-test (ns = not significant, ***p≤0.001).

## Materials and Methods

### hiPSC lines

#### Patient sample collection and reprogramming for iPSC line

Patient fibroblasts were retrieved under approval of Local Ethics Committee Radboud UMC (Centrale Commisie Mensgebonden Onderzoek, CMO-light, protocol number 2015-1543). The iPSC lines were newly generated in the framework of this study. Fibroblasts were reprogrammed with four factors (Oct4, Sox2, Klf4, cMyc) by using retroviral vectors, cultured in Essential 8 Flex Medium (A2858501, Life Technologies) and passaged weekly using 0.5 mM EDTA. Pluripotency was determined using quantitative PCR (qPCR) and immunofluorescence. Presence of the pathogenic variants was confirmed by Sanger Sequencing (Fig S1A-B).

#### Generation of isogenic variant iPSC lines

A healthy donor hiPSC line, LUMC0099iCTRL04 (RRID:CVCL_UK77), was used to create isogenic pairs for the *USH2A* and *COCH* patient variants: LUMC99-04 COCH c.151C>T and LUMC99-04 USH2A c.7595-2144A>G.

##### CRISPR-Cas9-mediated gene-editing

Gene editing of LUMC0099iCTRL04 was performed using the NEON Transfection System (Invitrogen). Cells were electroporated with a Cas9-RNP complex targeting *USH2A* or *COCH* and a ssODN repair template (IDT). The Cas9-RNP complex was assembled by incubating Cas9 protein (spCas9, Geijsen lab LUMC) with a sgRNA (IDT), following the manufacturer’s protocol. For *USH2A,* two sgRNAs were used (5’- TTAAAGATGATCTCTTATCT-3’ and 5’- CTCTTATCTTGGGAAAGGAG-3’) with an ssODN (5’- CTGGCTTTTAAGGGGGAAACAAATCATGAAATTGAAATTGAACACCTCTCCTTTCCCAAGGTAAGAGATCATCTTTAAGAAAAGGCTGTGTATTGTGGGGGTTTGAAGTGCAAGTTCATC-3’). For *COCH*, one sgRNA was used (5’- CCTCAAGAGGGCAGCCCCCT-3’) with an ssODN (5’- GCTATCACATGTTTTACCAGAGGCTTGGACATCAGGAAAGAGAAAGCAGATGTCCTCTGCTCAGGGGGCTGCCCTCTTGAGGAATTCTCTGTGTATGGGAACATAGTATATGCTTCTGTA-3’). After electroporation cells were plated on Synthemax II-SC-coated plates (0.45 μg/cm^2^) in mTesR-plus medium (STEMCELL Technologies) without antibiotics, supplemented with CloneR^TM^2 (STEMCELL Technologies, #100-0691) and grown at 37°C and 5% CO_2_.

###### Single cell cloning

Following recovery clonal lines were derived by limiting dilution. Cells were harvested as single-cells using Accutase dissociation reagent, strained using a 40 µm strainer, counted (Luna Automated fluorescent Cell counter) and 1000 cells were plated on a 10cm Synthemax II-SC-coated dish in mTeSR Plus with CloneR^TM^2 (100-0691, STEMCELL Technologies). Single cells were allowed to grow out in ∼10 days where medium was changed every other day, before clones were picked and/or passaged. Genomic DNA was isolated using QuickExtract DNA Extraction Solution (Lucigen, #QE0905T), and the target region was amplified by PCR (Terra PCR Direct Polymerase Mix, TaKaRa) using the following primers; for *USH2A* 5’- GTTGCAGGCCAGTTGATTTG-3’ and 5’- CTGTGAGCATCTTCAGGGAAA-3’; for *COCH* 5’- AATCTGGAATGGTATGGAAGGG-3’ and 5’- AGATGGGTAAAGCAGGAAAGG-3’. PCR products were digested using StyI restriction enzyme and Bpu10I restriction enzyme (ThermoFisher) at 37°C for *USH2A* and *COCH,* respectively. Restriction patterns were analyzed by agarose gel. Candidate clones were validated by Sanger sequencing (Leiden Genome Technology Centre using the ABI3730xl system). The edited iPSC lines were re-characterized and met all quality control criteria, including maintenance of pluripotency and genomic integrity.

#### hiPSC culture

All iPSCs lines used in this study were cultured on six-wells plates coated with vitronectin (A14700, Gibco, concentration 0.5 µg/ml) and maintained in mTesR^TM^ Plus Basal medium (100-0276, STEMCELL Technologies) with normocin (ANT-NR-1, Invivogen, concentration 100 µg/ml). Medium was refreshed every other day. Colonies were closely monitored for differentiation and passaged every 4-6 days (∼80% confluence) as small clusters using 0.5 mM EDTA (15575020, Invitrogen) in dPBS (14190250, Gibco).

### Generation of IEO from hiPSC cells

hiPSCs were differentiated into inner ear organoids using previously published protocols (20). Briefly, hiPSCs at 75% confluency were dissociated with accutase (A11105-01, Gibco) into single cells and resuspended in mTeSR with a 50 nM Chroman-1 (HY-15392, MCE), 5 µM Emricasan (HY-10396, MCE), 1x Polyamine (P8483, Sigma-Aldrich) and 700 nM Trans-ISRIB (5284, Tocris). Then cells were replated into a Nunclon Sphera 96-well round bottom plate (174929, Thermo Scientific) at a final cell concentration of 25,000 cells/ml to aggregate in embryoid bodies.

After two days, on day 0 of differentiation, the aggregates were collected using a W-O p1000 tip (17014297, Rainin), washed three times with Essential 6 medium (E6, A1516401, Gibco) to remove the mTeSR medium, and individually transferred with a W-O p200 tip (17014294, Rainin) into a 96-well plate in 100 µl E6 medium containing normocin (Ant-nr-1, Invivogen), 2% Matrigel GFR (354230, Corning), 10 µM SB431542 (04-0010-05, Stemgent), 4 ng/ml recombinant human FGF-basic (bFGF, 100-18B-100, Peprotech) and recombinant human BMP-4 (314-BPE, R&D). BMP4 concentrations ranged from 2,5 to 15 ng/ml and titrations were performed to reach optimal differentiations. On day 3 of differentiation, 25 µl E6 medium containing normocin, 1 µM LDN-193189 (final concentration 200 nM, 04-0074-02, Stemgent) and 250 ng/ml bFGF (final concentration 50 ng/ml) was added to the aggregates. On day 6 of differentiation, 75 µl E6 medium containing normocin was added. On day 8 of differentiation, using a multi-channel pipette, 100 µl medium was carefully removed, remaining 100 µl medium in each well. Into each well 100 µl fresh E6 medium containing normocin and 6 µM CHIR99021 (final concentration 3 µM, hereafter CHIR, 04-0004-02, Stemgent) was added. On day 10 of differentiation, again, 100 µl medium was carefully removed and 100 µl fresh E6 medium containing normocin and 3 µM CHIR was added.

On day 12 of differentiation, the aggregates were collected using a W-O p1000 tip and washed three times with Advanced DMEM/F12 medium (12634010, Gibco). Aggregates were then collected using a W-O p1000 tip and individually placed in a Nunclon Sphera 24-well plate (174930, Thermo Scientific) in 500 µl Organoid Maturation medium (hereafter referred to as OM) supplemented with 1% Matrigel and 3 µM CHIR. OM consisted of a 1:1 mix of Advanced DMEM/F12 and Neurobasal medium (10888022, Gibco) with the addition of 1x GlutaMax (35050061, Gibco), 0.5x B-27 supplement (minus Vit A, 12587-010, Gibco), 0.5x N-2 supplement (17502-048, Gibco), 0.1 mM 2-Mercaptoethanol (21-985-023, Thermo Scientific) and 100 µg/ml normocin (ant-nr-1, Invivogen). Plates were incubated on a rocker from day 12 until the end of differentiation. On day 15 of differentiation, 250 µl of spent medium was removed and 250 µl fresh OM supplemented with 1% Matrigel and 3 µM CHIR was added. On day 18 of differentiation, 250 µl of spent medium was removed and 250 µl fresh OM was added. Hereafter, a half medium change was performed every 2-3 days with a full medium change once per week. Based on the organoid size and medium consumption rate the total medium volume was increased to 1.5 ml per well to provide proper nutrition. Plates were incubated in a 37°C incubator with 5% CO_2_ throughout the differentiation process.

### Human adult inner ear sample collection

Human adult vestibular samples were collected from patients undergoing translabyrinthine surgery for vestibular schwannomas. The macula of the utricle was carefully excised by the surgeon. Tissue samples were immediately transported in OM medium with fungin (ant-fn-1, Invivogen) on ice. Under a Leica M205C dissecting microscope, debris and the otoconia were removed. The collection of human vestibular samples during translabyrinthine vestibular schwannoma surgery was approved by the Medical Research Ethics Committee of Leiden University Medical Center (protocol registration number BB23.001) and obtained after written informed consent of the donor.

### Anti-sense oligonucleotide experiments

#### ASOs design and synthesis

ASO sequences were previously designed (32,46). The ASOs were purchased from Eurogentec (Liège, Belgium) and SynOligo (Morrisville, NC, US). As controls, a sense ASO (SON) was acquired for the *USH2A-*targeting ASO, and a scrambled-sequence ASO (SCR) was acquired for the *COCH*-targeting ASO, as described in the previous publications (32,46). Upon arrival, the ASOs were dissolved in PBS to a 1 mM stock solution, aliquoted, and stored in a -80 °C freezer. All ASO sequences used in this study are presented in Table 1.

#### ASO delivery in inner ear organoids

To allow access of ASOs to disease-relevant cell populations, day 75 IEOs were sectioned into 200 µm slices with a vibratome (VT1200S, Leica), as previously described (57). Organoids were placed in a cryomold, liquid was removed and a 4% agarose solution (A0701-25G, Sigma-Aldrich) in PBS was added. Air bubbles were removed using a needle and the block was put on ice to solidify. Then, the agarose block containing the tissue was trimmed and attached to the vibratome tissue holder slide using Histoacryl (1050071, B.Braun). The vibratome cooling chamber was filled with ice. Organoids were sliced at a 0.2 mm/s rate, an amplitude of 1 mm and a slice thickness of 200 µm, using a razor blade (Astra). The vibratome slices were collected in ice-cold PBS. After slicing, the vibratome slices were transferred from the collection chamber into medium-containing 12-well plates and placed in a 37°C incubator with 5% CO2. Vibratome slices were checked under a Leica IC90 E microscope for integrity and to confirm otic identity. Vibratome slices were left to recover for two days before proceeding with ASO experiments.

On day 0 of the ASO IEO experiments, ten vibratome slices were placed together in 500 µl OM with fungin (ant-fn-1, Invivogen) in a Nunclon Sphera 24-well plate (174930, Thermo Scientific). ASOs were dissolved in OM to the desired concentration and added to the respective conditions. For *USH2A* experiments, a single dose of ASOs was added on day 0. Half medium changes were performed every 3 days. Samples were collected after 28 days for downstream analysis. For *COCH* experiments, a repeated dosing regime was followed by adding fresh ASOs on day 0, 3, 7 and 11. Samples were collected after 12 days for downstream analysis. For AlexaFluor 488-labeled ASO experiments, during the culture period, pictures were taken on days when medium changes were performed using a EVOS M5000 microscope.

#### ASO delivery in human adult inner ear tissue

Culture conditions of the human adult inner ear tissue were adapted from a previous study (58). After microscopic evaluation, samples were transferred to a 48-well plate in 200 µl OM medium with fungin (ant-fn-1, Invivogen) containing 10 µM ASO (Table 1). The surrounding wells were filled with dPBS (14190250, Gibco). Samples were incubated at 37°C and 5% CO_2_. After 48h, a half medium change was performed. On day 4, the samples were fixed in 4% paraformaldehyde overnight and processed for downstream analysis.

### Gene expression analyses

#### Gene expression analysis in the prenatal inner ear atlas

*USH2A* and *COCH* expression were examined using our previously published transcriptomic atlases of human prenatal inner ear development (HIEDRA; (41)) and IEOs (19). Expression of each gene was visualized across all annotated cell clusters using density plots generated with Nebulosa (59). Expression was additionally summarized across clusters using a dot plot generated with ggplot2 (60). Cluster identities were assigned as described in the original atlas.

#### RNA extraction

RNA was extracted from IEOs using the Monarch kit (T2110 or T2010). Samples were collected in StabiLyse DNA/RNA buffer with nuclease-free water in a 1:1 ratio. On the day of RNA isolation, samples were equilibrated to room temperature. Mechanical homogenization was performed for 30 sec with a tissue grinder, samples were placed on ice for at least 2 min. For every 350 µl StaibiLyse DNA/RNA buffer and sample mixture, 15 µl Proteinase K was added. Samples were vortexed briefly and incubated for 30 min at 55°C. After Proteinase K incubation, an equal volume of StabiLyse DNA/RNA buffer was added and samples were briefly vortexed again. The samples were loaded on a gDNA removal column fitted with a collection tube and centrifuged for 1 min to remove most of the gDNA. An equal volume of ethanol (>95%) was added to the flow-through and mixed by pipetting. The mixture was transferred to a RNA purification column fitted with a collection tube and centrifuged for 1 min. Flow-though was discarded and 700 µl Monarch Buffer WZ was added to the column before centrifuging for 1 min. A on-column DNase I treatment was performed for 15 mins at room temperature. The column was washed with 500 µl Monarch Buffer BX, spin for 1 min, followed by two washes with 500 µl Monarch Buffer WZ for 1 min and 2 mins. Finally, RNA was eluted by adding 20-100 µl nuclease-free water to the column and spinning for 30 sec. All centrifugation steps were perfomed at 16,000 x *g*. RNA concentration was measured using NanoDrop One (Thermo Fisher Scientific) and samples were stored at -80°C.

#### cDNA synthesis

For *COCH* gene expression analysis, cDNA was generated according to the iScript manual (1708891, iScript kit, Bio-Rad). A reaction mixture was made consisting of iScript reaction mix, the iScript reverse transcriptase, nuclease-free water and the RNA template (200-500 ng). Thermal cycling (Bio-Rad S1000 Thermal Cycler) took place by priming for 5 min at 25°C, reverse transcription for 20 min at 46°C, reverse transcription inactivation for 1 min at 95°C, and cooled down to 4°C. cDNA was stored at -80°C.

For *USH2A* gene expression analysis, cDNA was synthesized using the SuperScript IV First-Strand Synthesis Kit (18091150, Thermo Fisher Scientific). First, a RNA-primer mix was made by combining 50 µM oligo d(T)_20_ primers, 10 mM dNTP mix, nuclease-free water and the RNA template (200-500 ng). The RNA-primer mix was heated for 5 min at 65°C and incubated on ice for at least 1 min. In a separate reaction tube, the reverse transcriptase reaction mixture was made by combining 5x SuperScript IV (SSIV) buffer, 100 mM DTT, ribonuclease inhibitor and SuperScript IV Reverse Transcriptase (200 U/µl). The reverse transcriptase reaction mixture was added to the RNA-primer mix and incubated for 50 min at 55°C. The reaction was inactivated by incubating for 10 min at 80°C. The cDNA was then kept on ice and transferred to -80°C for long term storage. RNA samples derived from ASO experiments were heated for 1 min at 95°C prior to cDNA synthesis.

#### Reverse transcription PCR (RT-PCR) for *USH2A*

RT-PCR products for *USH2A* were amplified using the Q5 High-fidelity polymerase kit (M0491S, New England Biolabs). Reaction mixture was generated by combining the Q5 high-fidelity 2x master mix (1x), 10 µM forward primer 5’- GCTCTCCCAGATACCAACTCC-3’(0.5 µM), 10 µM reverse primer 5’- GAGGGTCAGGCATGTGAATC-3’(0.5 µM) and 500-1000 ng template RNA. Thermal cycling (Bio-Rad S1000 Thermal Cycler) occurred with a touchdown protocol starting with an initial denaturation for 30 sec at 98°C, 10 cycles with each cycle consisting of 10 sec at 98°C, 20 sec at 62.5-57.5°C, 1 min at 72°C, followed by 23 cycles with each cycle consisting of 10 sec at 98°C, 20 sec at 57.5°C, 1 min at 72°C, the protocol was ended with a final extension for 5 min at 72°C. The obtained PCR products were then loaded onto a 1.5% agarose gel for visualization.

#### TaqMan assay for *COCH* transcript levels

A previously designed allele-specific TaqMan assay for *COCH* variant (c.151C>T, p.Pro51Ser) was used (32). The TaqMan assay used primers 5′-GGACATCAGGAAAGAGAAAGCAGAT-3′ and 5′-CCCATACACAGAGAATTCCTCAAGAG-3′, a wild-type allele-specific HEX-labeled probe 5′-CCCCCTGGGCAGAG-3′ and a mutant allele-specific FAM-labeled probe 5′-CCCCCTGAGCAGAG-3′. Assay specificity was validated using synthetic WT and mutant gBlock templates, showing ∼2% cross-reactivity between probes, and 3-4% in real biological samples (Figure S3A). Reaction mixtures were set up by combining 2x TaqMan Fast Advanced Master Mix (4444556, Thermo Fisher Scientific), 40x custom TaqMan SNP Genotyping Assay (4332077, Assay ID ANMFYDW, Assay name P51S, Thermo Fisher Scientific), nuclease-free water and 500 ng cDNA. Reaction was run on a real-time PCR system (Bio-Rad CFX Opus 384) thermal cycling program starting at 50°C for 2 min, then 95°C for 20 s, followed by 40 cycles of 95°C for 3 sec and 60°C for 30 sec. The CFX Maestro (Bio-Rad) software was used to analyze the data. Samples were measured in triplicates.

#### Quantitative PCR (qPCR)

For other gene expression analyses, a reaction mix was prepared of 10 µM forward and 10 µM reverse primers, iQ SYBR Green 2x master mix (1708882, Bio-Rad) and 5x diluted cDNA. A list of primers can be found in Table 2. A real-time PCR system (Bio-Rad CFX Opus 384) thermal cycling program was run at 95°C for 3 min, followed by 60°C for 30 sec and 72°C for 40 sec, in a total of 40 cycles. After which samples were kept at 4°C. The CFX Maestro (Bio-Rad) software was used to analyze the data. Samples were measured in triplicates.

**Table 2.**
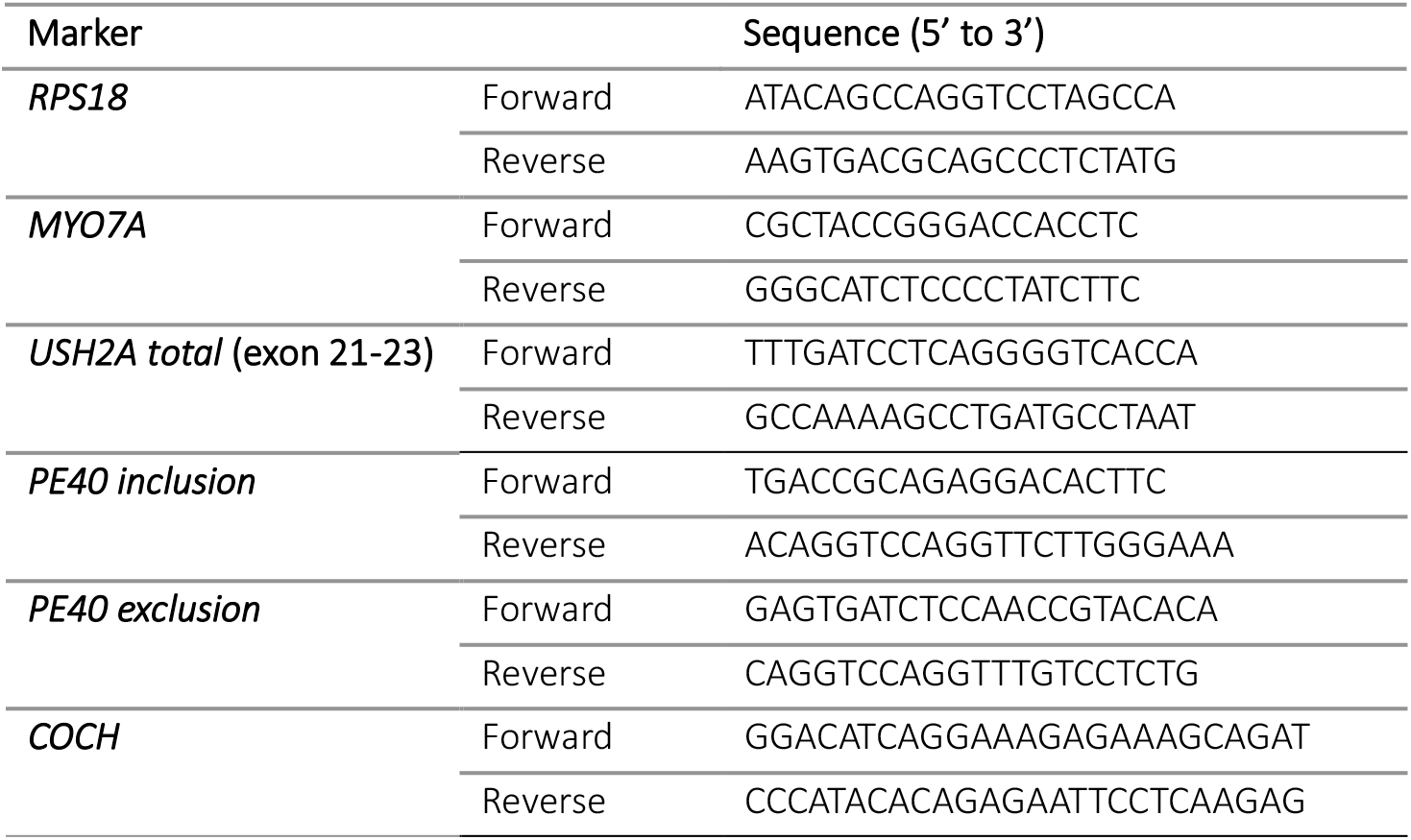
List of primers.

#### Statistical analysis

Statistical analyses were performed using GraphPad Prism (version 10.2.3). A *P*-value of ≤0.05 was considered statistically significant for all analyses. The specific statistical tests used for each analysis are indicated in the corresponding figure captions.

### Tissue processing and immunohistochemistry

#### Immunohistochemistry and image acquisition

For immunohistochemical analysis, samples were first embedded in Histogel (HG-4000-012, Epredia) and subsequently processed and paraffin-embedded. Paraffin blocks were sectioned into 5 μm slices using a rotary microtome (HM355S, Thermo Fisher Scientific). Prior to staining, sections were deparaffinized in xylene and rehydrated through a graded ethanol series of decreasing concentration, followed by rinsing with deionized water. One section from every set of ten slides was selected for hematoxylin and eosin (H&E) staining using Hematoxylin (40859001, Klinipath) and Eosin (AB246823, Abcam).

For immunofluorescent staining, antigen retrieval was carried out in 10 mM sodium citrate buffer (pH 6.0; S1804-50G, Sigma-Aldrich) at 97°C for 12 min. Sections were then washed with PBS containing 0.05% Tween-20 (H5152, Promega) and incubated for 30 min in blocking solution composed of PBS, 0.05% Tween-20, and 5% bovine serum albumin. Primary antibodies (Table 3) were diluted in blocking buffer and applied overnight at 4°C in a humidified chamber.

**Table 3:** List of primary and secondary antibodies used in this study.

| Primary antibodies |  |  |  |  |  |  |
| --- | --- | --- | --- | --- | --- | --- |
| Marker | Host | Clonality | Company | Cat. No. | Dilution | RRID |
| MYO7A | Mouse | Monoclonal | DSHB | 138-1-s | 1:50 | AB_2282417 |
| MYO7A | Rabbit | Polyclonal | Proteus | 25-6790 | 1:500 | AB_10015251 |
| SOX10 | Goat | Polyclonal | ThermoFisher Scientific | PA5-47001 | 1:100 | AB_2608449 |
| CDH1 | Mouse | Polyclonal | BD Biosciences | 610182 | 1:20 | AB_397581 |
| Tubelin beta 3 class III (TUBB3) | Mouse | Monoclonal | R&D systems | MAB1195 | 1:100 | AB_357520 |
| COCH | Rabbit | Polyclonal | LSBio | LS-C334796 | 1:100 | AB_3752095 |
| USH2A | Mouse | Monoclonal | Bio-Tem | Custom made against the C-terminus | 1:100 | n/a |
| ASO PS backbone Moddetect Ps05 | Mouse | Monoclonal | Rockland | 200-301-MV1 | 1:100 | AB_3752096 |
| Secondary antibodies |  |  |  |  |  |  |
| Host | Antibody against | Alexa Fluor | Fisher Scientific Catalog No. | RRID |  |  |
| Donkey | Mouse IgG (H+L) | 488 | A21202 | AB_141607 |  |  |
| Donkey | Rabbit IgG (H+L) | 594 | A21207 | AB_141637 |  |  |
| Donkey | Mouse IgG (H+L) | 594 | A21203 | AB_2535789 |  |  |
| Donkey | Goat IgG (H+L) | 680 | A21084 | AB_141494 |  |  |
| Donkey | Rabbit IgG (H+L) | 680 | A10043 | AB_2534018 |  |  |
| Donkey | Mouse IgG (H+L) | 680 | A10038 | AB_11180593 |  |  |

Following three washes of 5 min each in washing buffer, sections underwent an additional 10 min blocking step before incubation with secondary antibodies (1:500 dilution; Table 3) and DAPI (1:1000 dilution) for 60 min at room temperature. After a final washing step, coverslips were mounted using ProLong™ Gold Antifade Mountant (P36934, Invitrogen).

Imaging was performed on a Leica SP8 WLL1 inverted equipped with Leica objectives (HC PL APO CS2 20x/0.75 DRY, HC PL APO CS2 40x/1.30 OIL, or HC PL APO CS2 63x/1.40 OIL). Image acquisition was conducted using LAS X software (Leica, version 3.7.6.25997). The IHC images in Figure S3B were taken on a Zeiss LSM900 Airyscan upright microscope using Plan-Apochromat 63x/1.40 Oil DIC objective, for which the image acquisition was conducted using Zeiss ZEN 3.0 software. All presented images are optical z-stack sections obtained from paraffin-embedded tissues. Brightness and contrast adjustments were performed with Fiji (ImageJ, version 1.54t).

#### BaseScope and miRNAscope

For the BaseScope and miRNAscope experiments samples were embedded into paraffin blocks, 5µm thick sections were cut and mounted on Superfrost Plus glass slides (12-550-15, Fisher Scientific).

BaseScope was performed according to the BaseScope^TM^ Duplex Detection Reagent Kit manual (ACD, 323800-USM). Probes were designed against USH2A-WT (HRP, Green) and USH2A-Mutant (AP, Fast Red) and obtained from ACD/Biotechne. Slides were prebaked at 60°C for 1 h and then FFPE sections were deparaffinized. Samples were treated with RNAscope Hydrogen Peroxide at RT for 10 min. Manual target retrieval was performed by submerging the slides in mildly boiling RNAscope Target Retrieval Reagent solution for 15 min, followed by protease treatment with miRNAscope Protease III at 40°C for 30 min. Probe hybridization was performed at 40°C for 2 h, followed by washing and incubation with the 8 amplification steps, before Fast Red signal detection at RT for 10 min, followed by amplification steps 9-12, and detection of the green signal at RT for 10 min. Gill’s hematoxylin (GHS132, Sigma-Aldrich) was used as a counterstain for nuclei detection. All steps at 40°C were performed in a HybEZ^TM^ hybridization system. Slides were dried and then mounted with VectaMount permanent mounting medium (H-5000, Vector labs). To obtain digital whole slides, stained slides were scanned with a 2DHISTECH P250 slidescanner with a 20x objective.

miRNAscope was performed according to the miRNAscope^TM^ HD (RED) assay manual (ACD, UM324510). A target probe was designed against the PE40-ASO (Fast red) and obtained from ACD/Biotechne. Slides were prebaked at 60°C for 1 h and then FFPE sections were deparaffinized. Samples were treated with RNAscope Hydrogen Peroxide at RT for 10 min. Manual target retrieval was performed by submerging the slides in mildly boiling RNAscope Target Retrieval Reagent solution for 15 min, followed by protease treatment with miRNAscope Protease III at 40°C for 30 min. Probe hybridization was performed at 40°C for 2 h, followed by repeated washing and incubation with the 6 amplification steps, before signal detection at RT for 10 min. Gill’s hematoxylin (GHS132, Sigma-Aldrich) was used as a counterstain for nuclei detection. All steps at 40°C were performed in a HybEZ^TM^ hybridization system. Slides were dried and then mounted with EcoMount permanent mounting medium (EM897L, Biocare). To obtain digital whole slides, stained slides were scanned with a 2DHISTECH P250 slidescanner with a 20x objective.

#### Whole-mount clearing, staining and image acquisition

Whole-mount immunostaining and clearing were performed on AlexaFluor 488-labeled PE4-ASO treated patient-derived inner ear organoids and ASO-treated adult human inner ear tissue using a modified quantitative clearing-enhanced 3D (qCe3D) protocol (61). Samples were fixed in 4% paraformaldehyde (PFA), followed by permeabilization and blocking at 37°C for 8 h with gentle agitation using a blocking buffer containing 2% bovine serum albumin (A3059, Sigma-Aldrich) and 0.3% Triton X-100 (T-9284, Sigma-Aldrich). Tissues were incubated with primary antibodies (Table 3) diluted in staining buffer for 2 days at 37°C with gentle agitation to facilitate antibody penetration throughout the sample. Following washing with washing buffer (1x PBS supplemented with 0.3% Triton X-100 and 0.5% 1-thioglycerol), samples were incubated with fluorophore-conjugated secondary antibodies and DAPI (Invitrogen, D3571) for 1.5 days under the same conditions. After immunolabeling, tissues were washed thoroughly and subjected to Ce3D optical clearing. The Ce3D clearing solution consisted of 40 % *N*-methylacetamide (M26305-500G, Sigma-Aldrich), Histodenz (D-2158, Sigma-Aldrich), 1-Thioglycerol (M1753, Sigma-Aldrich) and Triton X-100 (T-9284, Sigma-Aldrich) and was prepared according to the instructions in Lee et al (2022). Samples were incubated in Ce3D clearing solution o/n until transparency was achieved and subsequently mounted in fresh clearing solution for imaging. AlexaFluor 488-labeled PE40-ASO treated patient-derived inner ear organoids samples were imaged using a spinning-disk confocal microscope, Andor DragonFly 500 inverted equipped with (objectives HCX PL APO 40x/1.30 OIL and HCX PL APO 63x/1.40-0.60 OIL). ASO-treated adult human inner ear tissue was imaged on a Leica SP8 WLL1 inverted equipped with Leica objectives (HC PL APO CS2 20x/0.75 DRY, HC PL APO CS2 40x/1.30 OIL, or HC PL APO CS2 63x/1.40 OIL). Image acquisition was conducted using LAS X software (Leica, version 3.7.6.25997). Image acquisition Z-stack images were acquired throughout the tissue volume. Brightness and contrast adjustments were performed with Fiji (ImageJ, version 1.54t).

