## Supplementary Figures for "Patient-Derived Inner Ear Organoids as a Disease Modeling and Therapy Validation Platform For Hereditary Inner Ear Disorders"

#### Figure S1

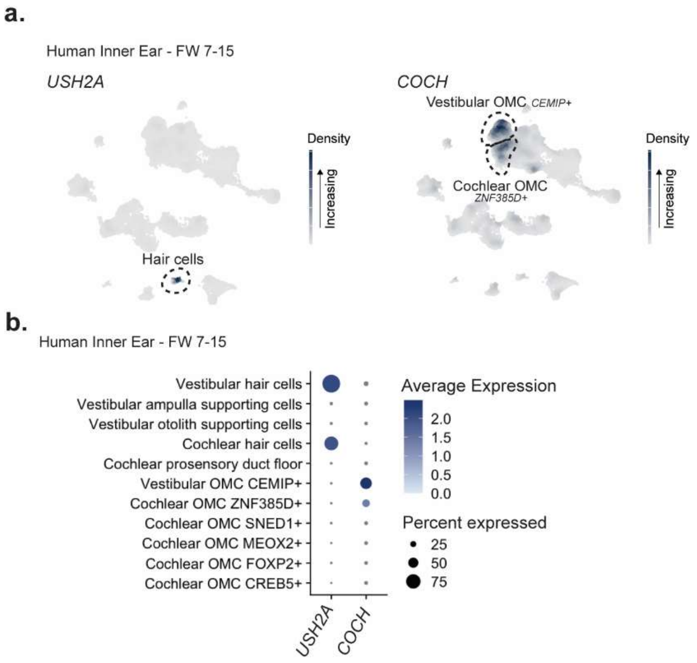


**Figure S1** Expression of *USH2A* and *COCH* in human fetal inner ear.

a. Feature plots showing gene expression of *USH2A* and *COCH* in single-nuclei RNAseq data from fetal week 7-15, retrieved from Human Inner Ear Development single cell RNA Atlas (HIEDRA, (41)).

b. Dot plot showing *USH2A* and *COCH* gene expression levels in relevant inner ear cell types, indicating the average relative expression among expressing cells (dot color) and the percentage of cells expressing the gene within each annotated cell type (dot size).

Figure S2

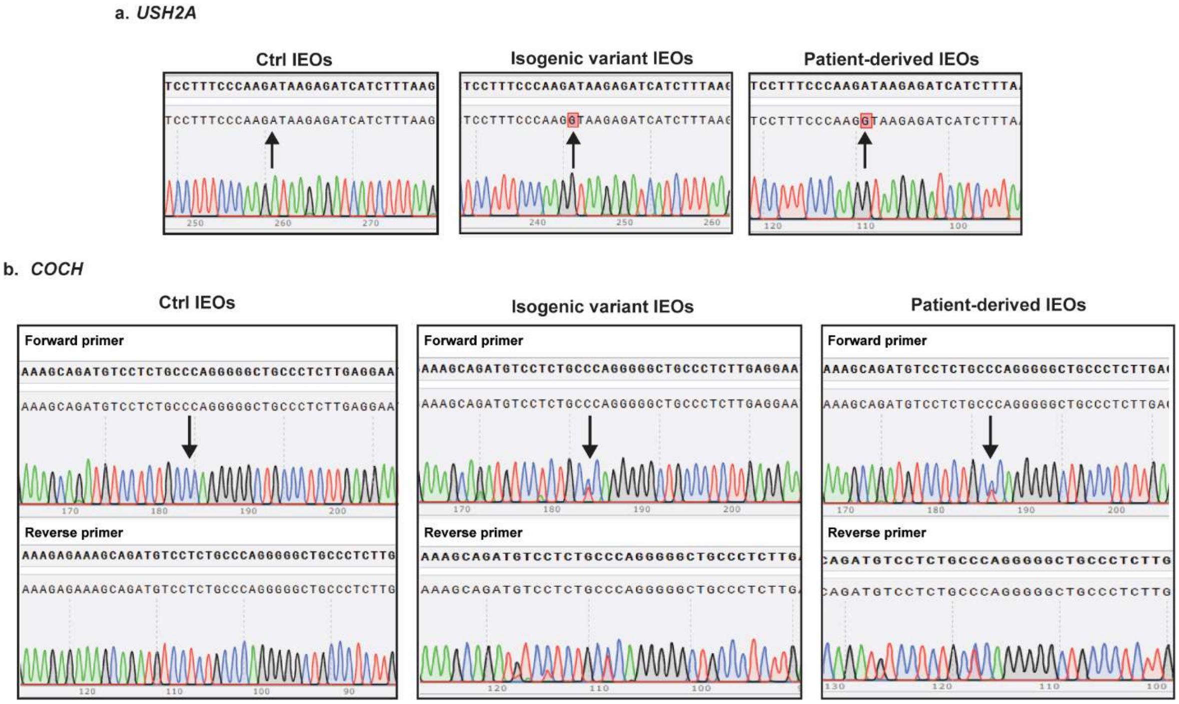


Figure S2 – Confirmation of genotypes with Sanger Sequencing

A. *USH2A* (c.7595-2144A>G, homozygous) in control, isogenic variant and *USH2A*-patient derived IEOs. Arrows indicate the site of the variant.
B. *COCH* (c.151C>T; p.Pro51Ser, heterozygous) in control, isogenic variant and *COCH*- patient derived IEOs. Arrows indicate the site of the variant.

### Figure S3

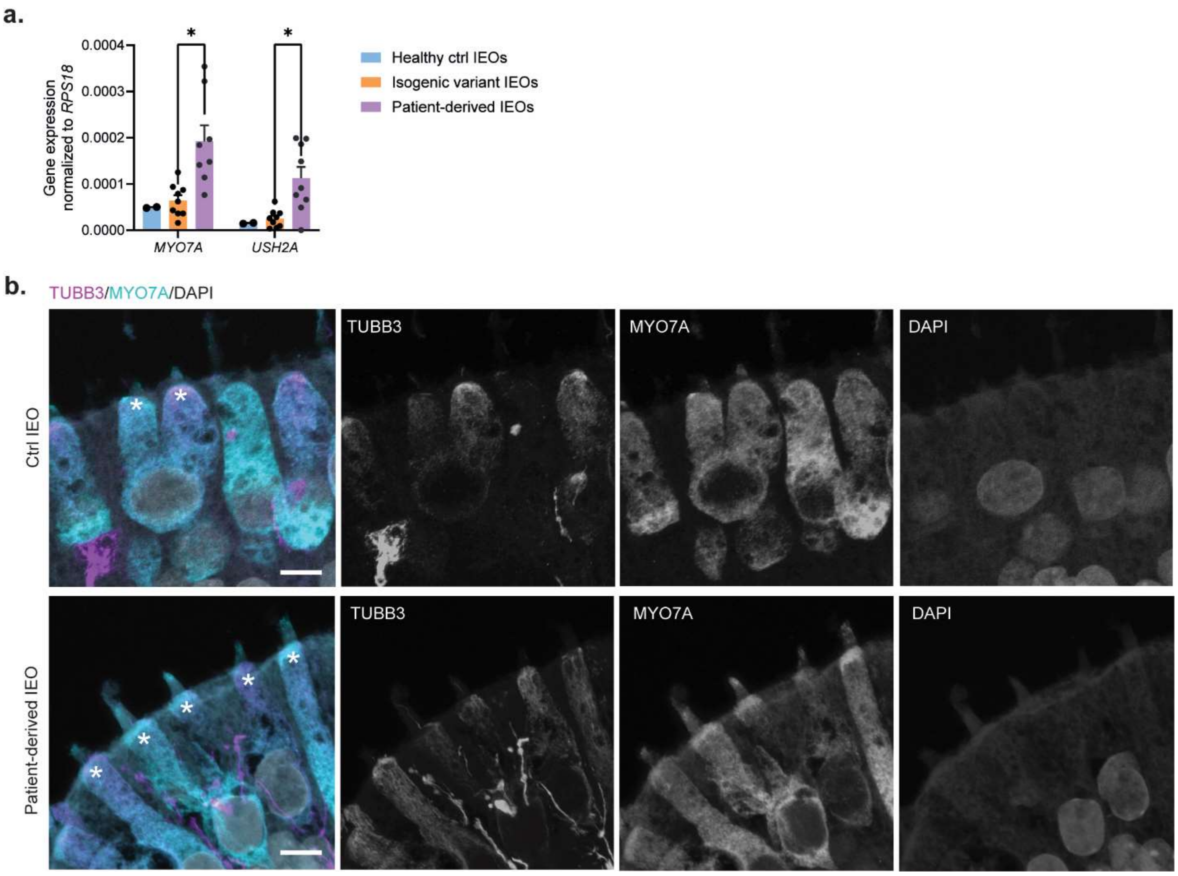


Figure S3 Additional information *USH2A*

a. Gene expression of *MYO7A* and *USH2A* in day 75 control (*n* = 2), isogenic variant (*n* = 9) and *USH2A* patient-derived (*n* = 9) inner ear organoids (IEOs), normalized to *RPS18* expression. Statistical test via multiple unpaired t-tests with Welch correction (*p≤0.001).

b. Representative panels showing immunofluorescence (IF) staining of day 75 IEOs derived from the control and *USH2A* patient-derived iPSCs, containing hair cells (MYO7A⁺, red) and neurons (TUBB3⁺, yellow). * indicate the base of the stereocilia bundle. Scale bars: 5 µm.

### Figure S4

**a.**

**TaqMan assay with gBlocks**
**Cross-reactivity**

| | 500 | 1000 | 5000 | 50000 | Healthy IEOs |
|---|---|---|---|---|---|
| % WT bleeding into mutant channel | 2,05 | 2,42 | 3367,206 | 3351,191 | 3,65 |
| % Mut bleeding into WT channel | 0,05 | 0,03 | 0,026 | 0,021 | |

**b.**

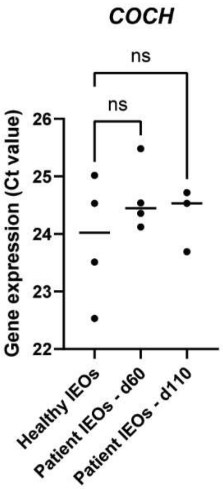


Figure S4 – Additional information for *COCH*

a. Cross-reactivity coefficients calculated from TaqMan assay wild type (WT) and mutant *COCH* c.151C>T expression using gBlocks of wild type and mutant *COCH* in 500-50000 copies/µl. Cross-reactivity = 2^(-A) / 2^(-B) * 100, in which A = ‘Ct value_gBlock WT – Mut channel’ and B = ‘Ct value_gBlock WT – WT channel’.

b. Gene expression (Ct value) of the *COCH* gene upon normalized RNA input in day 75 control (*n* = 4), patient IEOs day 60 (*n* = 4) and patient IEOs day 110 (*n* = 3). Statistical test via ordinary one-way ANOVA (ns = non-significant).

### Figure S5

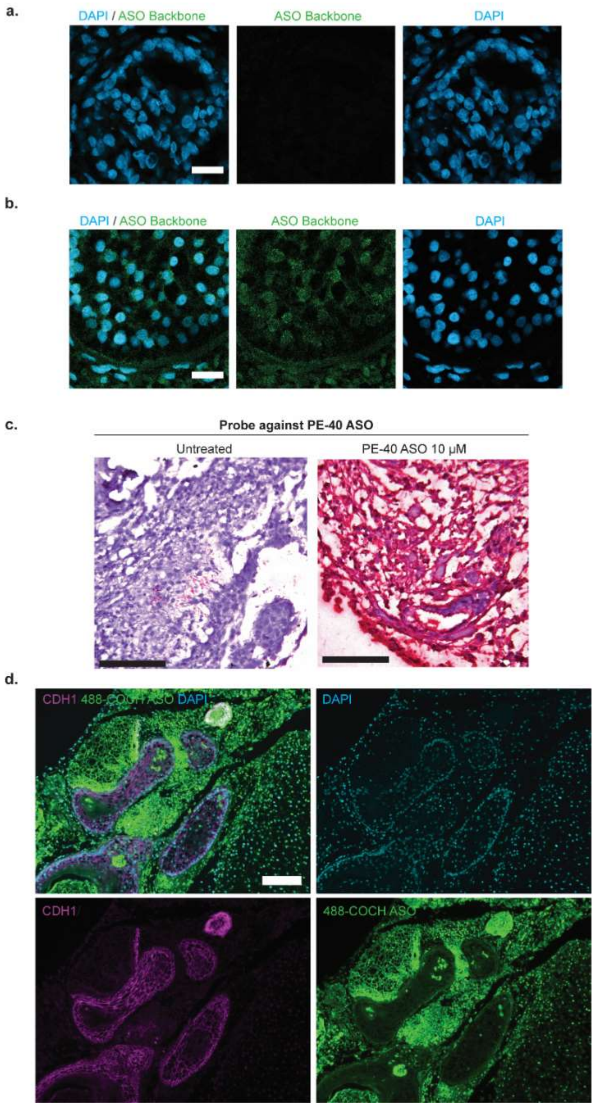


Figure S5 – Additional ASO uptake experiments.

a. Representative panels showing negative control immunofluorescence staining of the localization of the ASO backbone (green) in off-target tissues in patient-derived IEOs after 28 days treatment with 10 μM splice-switching ASO targeting pseudoexon-containing *USH2A* variant. Scale bar: 20 μm.
b. Representative panels showing immunofluorescence staining of the localization of the ASO backbone (green) in off-target tissues in patient-derived IEOs after 28 days treatment with 10 μM splice-switching ASO targeting pseudoexon-containing *USH2A* variant. Scale bar: 20 μm.
c. Probe against PE40 *USH2A* ASO shows localization of PE40-ASO throughout the IEO treated with 10 μM splice-switching ASO. Scale bar: 100 μm.

d. Representative panels showing IF staining of the localization of the AF488-labeled gapmer ASO targeting COCH, showing that the labeled ASO (green) does not localize in epithelial cells ($CDH1^+$, magenta). Scale bar: 100 µm.

### Figure S6

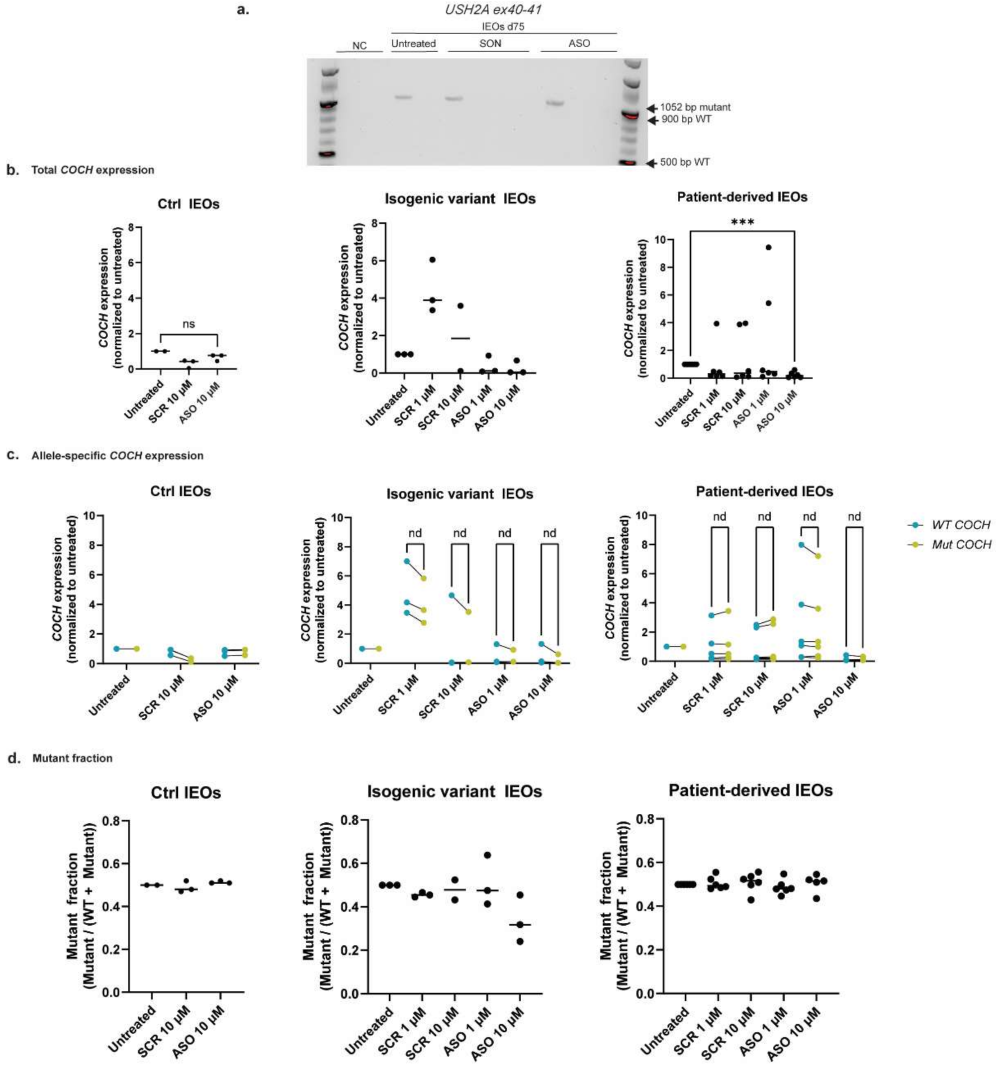


Figure S6 Total *COCH* and allele-specific *COCH* expression in ctrl IEOs, isogenic variant IEOs and patient-derived IEOs.

a. Absence of wild type transcript in day 75 *USH2A*-patient-derived IEOs treated with PE40-ASO for 12 days, as shown by reverse transcriptase (RT)-PCR targeting *USH2A* ex40-41. NC indicates a negative control sample, SON indicates a nonbinding sense oligonucleotide.

b. Total *COCH* expression in control IEOS ($n$ = 3), isogenic variant IEOs ($n$ = 3) and patient-derived IEOs ($n$ = 6) as shown by quantitative, normalized to the untreated sample. Statistical test via Welch's t-test (ns = not significant, ***p≤0.001).

c. Allele-specific wild-type (WT, blue) and mutant (Mut, green) *COCH* expression for control IEOs ($n$ = 3), isogenic variant IEOs ($n$ = 3) and patient-derived IEOs ($n$ = 6) in untreated and ASO conditions, normalized to untreated samples. Statistical test via multiple paired t-tests (nd = non discovery).

d. Mutant fraction was calculated for the wild-type and mutant *COCH* expression as mutant / (WT + Mutant). Statistical test via Welch's t-test (ns = not significant, ***$p \leq 0.001$).
